# PSGL-1 directs early TCR signaling to repress metabolism and promote T cell exhaustion by modulating the TCF-1/TOX axis in CD8^+^ T cells

**DOI:** 10.1101/2022.01.24.477602

**Authors:** Jennifer L. Hope, Dennis C. Otero, Eun-Ah Bae, Christopher J. Stairiker, Ashley B. Palete, Hannah A. Faso, Monique L. Henriquez, Hyungseok Seo, Xue Lei, Eric S. Wang, Roberto Tinoco, Alexandre Rosa Campos, Jun Yin, Peter D. Adams, Anjana Rao, Linda M. Bradley

**Author notes:** Corresponding author: Linda M. Bradley, **Corresponding Author Email Address:**.

## Abstract

We previously identified the adhesion molecule PSGL-1 as a T cell intrinsic immune checkpoint regulator of T cell exhaustion. Here we show that the ability of PSGL-1 to restrain TCR sginaling correlates with decreased expression of the Zap70 inhibitor *Ubash3b* (Sts-1) in PSGL-1-deficient T cells. PSGL-1-deficency in T cells supports antitumor responses to a PD-1 blockade resistant melanoma wherein tumor-specific CD8^+^ T cells sustain an enhanced metabolic state, with an elevated metabolic gene signature that promotes increased glycolysis and glucose uptake to support effector functions. In models of chronic virus infection and cancer, this outcome was associated with CD8^+^ T cell stemness, as PSGL-1 deficient CD8^+^ T cells displayed increased TCF-1 and decreased TOX expression, a phenotype shown to be crucial for responsiveness to checkpoint inhibition. Further, we demonstrate that PSGL-1 signaling promotes development of exhaustion in human CD8^+^ T cells. Finally, pharmacologic blockade of PSGL-1 was sufficient to curtail T cell exhaustion and enhance functionality both with melanoma tumors and chronic LCMV infection, demonstrating that PSGL-1 represents a therapeutic target for immunotherapy for PD-1/PD-L1 resistant tumors.

---

The immune system, and in particular effector CD8^+^ T cells (T_EFF_), are vital in protecting against and clearing virally-infected cells and tumor cells. However, in the settings of chronic viral infection and cancer, multiple factors, including sustained exposure to antigen and the suppressive tumor microenvironment (TME), promote the development of T cell exhaustion. T cell exhaustion is characterized by a progressive decline in cytoxic function, loss of effector cytokine production, decreased proliferative capacity, and increased expression and co-expression of multiple inhibitory receptors including PD-1, CTLA-4, LAG3, and TIM-3^1^. The discovery of immune checkpoint blockade (ICB) monoclonal antibody therapy as a means to invigorate and promote anti-tumor immunity by preventing engagement of CTLA-4 and/or PD-1 on T cells is one of the greatest cancer therapeutic breakthroughs of the last decade with efficacy shown across many cancers including metastatic melanoma, Hodgkin’s lymphoma, and non-small cell lung carcinoma^2^. Despite dramatic success in some patients, particularly metastatic melanoma, the majority of patients are ultimately refractory to ICB due to acquired resistance to treatment. It is therefore essential to identify novel mechanisms regulating T cell exhaustion that could expand ICB’s applicability to a greater number of patients.

It is well established that adaptations in tumor cell metabolism supporting growth profoundly impact the TME. Only recently has it been recognized that within tumors, where infiltrating T cells (TILs) compete with cancer cells for nutrients, metabolic reprogramming of T cells has major consequences including dysfunction in antitumor T cell responses or limiting the rescue of exhausted T cells (T_EX_)^3,4^. Optimal TCR engagement together with co-stimulation initiates metabolic reprogramming of T cells from a resting state that utilizes oxidative phosphorylation (OXPHOS) to an activated, glycolytic state that supports T_EFF_ development and function via glucose uptake and metabolism utilizing the Akt/mTOR pathway^5^. Much less is known regarding the extent of TCR stimulation during prolonged antigen exposure in the TME. However, recent studies demonstrate that T_EX_ exhibit greatly inhibited TCR signaling with chronic LCMV infection^6^, which is also likely to occur in the TME. Decreased glucose availability results in attenuated glycolytic metabolism^7^, which is sufficient to dramatically reduce T cell signaling and effector gene expression in TILs ^8^ as well as upregulate PD-1^9^. However, competition for glucose in the TME does not fully account for T cell metabolic insufficiency underlying T cell dysfunction in cancer. Oxygen deprivation within the TME also impairs T cell activation and responses. Recent studies show that the loss of intra-mitochondrial respiration (OXPHOS) and biogenesis in TILs is also a driver of T_EX_^10^ and of lack of responsiveness to PD-1 blockade^11^. It is therefore imperative to identify mechanisms of checkpoint inhibition that can counterbalance the metabolic constraints of T_EX_ to promote T_EFF_ responses.

PSGL-1 (P-selectin glycoprotein ligand-1) is a cell-surface receptor found on most hematopoietic cells, including T cells^12,13^. This highly conserved protein is recognized for its capacity to mediate leukocyte migration upon binding to the selectin family members P, E, and L selectin^14^. Studies in humans and mice have identified that PSGL-1 also binds the chemokines CCL19 and CCL21^15^. More recently VISTA, a PD-L1 homologue, was shown to be a ligand of PSGL-1 under highly acidic conditions, which can occur in solid tumors^16^ as well as in lymph nodes^17^. Previously, we identified that genetic deletion of PSGL-1 prevented development of T_EX_ and supported viral clearance in the chronic LCMV model, LCMV Clone 13 (Cl13)^18^. In addition, PSGL-1 deficiency promoted growth control of an anti-PD-1 resistant melanoma tumor^18^. These outcomes were linked to increased production of cytokines and cytotoxic molecules, and to decreased expression of multiple inhibitory receptors by both CD4^+^ and CD8^+^ T cells. Further, we recently demonstrated that PSGL-1 deficiency during acute viral infection with LCMV Armstrong (Arm) promotes greater T_EFF_ and memory precursor T cell formation^19^, underscoring the important role PSGL-1 plays as a fundamental regulator of T cell responses as well as exhaustion.

In our current study, we investigated cellular and molecular mechanisms by which intrinsic PSGL-1 signaling promotes the development of T_EX_. Single-cell RNA-sequencing identified the differential regulation of genes associated with T cell metabolism and particularly glycolysis in PSGL-1 deficient tumor-specific CD8^+^ TILs. Upon activation, PSGL-1^*-/-*^ CD8^+^ T cells were more glycolytically active and exhibited increased glucose uptake. Because of these robust and rapid metabolic changes, we hypothesized that PSGL-1 regulates T cell receptor signaling. We now show localization of PSGL-1 with the TCR with co-ligation, and with PSGL-1 deficiency, increased sensitivity to TCR stimulation that was attributable to decreased expression of the ZAP-70 inhibitor, Sts-1^20^, and greater activation of the ERK1/2 and PI3K/Akt pathways. Notably, PSGL-1 deficiency in T cells led to an increased frequency of stem cell-like T cells (T_SC_) that express TCF-1 but low levels of TOX, which retain attributes of T_EFF_ and promote antitumor responses that are associated with responsiveness to ICB^21,22^. Our studies have consistently demonstrated a link between PSGL-1 signaling and upregulation of PD-1 on T cells. We now find that concomitant PSGL-1 signaling with repeated TCR stimulation of human T_EFF_ promotes functional exhaustion. Together, our data show that PSGL-1 signaling in CD8^+^ T cells attenuates TCR signaling and promotes the development of T_EX_ by inhibiting glycolysis. Importantly, blockade of PSGL-1 recapitulates PSGL-1 deficiency in both antitumor and antiviral responses, underscoring the translational potential of targeting PSGL-1 by ICB, alone or together with current immunotherapies.

## Results

### PSGL-1 restrains the magnitude and duration of TCR signaling

We previously discovered that the absence of PSGL-1 promoted enhanced effector CD4^+^ and CD8^+^ T cell development and decreased T cell exhaustion in mice infected with LCMV Cl13 or inoculated with either of the melanoma tumors, B16-OVA or YUMM1.5^18^. Subsequently, we found that PSGL-1 deficiency promotes greater T_EFF_ responses and formation of progenitor-like memory T cells after acute infection with LCMV Arm^19^. We therefore sought to determine how PSGL-1 expression modulates CD8^+^ T cell responses early after TCR stimulation. Naïve WT and PSGL-1^*-/-*^ OT-I CD8^+^ T cells were stimulated overnight (18 hours) with optimal and suboptimal concentrations of plate-bound anti-CD3 antibody and T cell activation was evaluated by flow cytometry. In response to TCR stimulation, PSGL-1^-/-^ CD8^+^ T cells displayed greater activation as indicated by increased expression of CD25, CD69, CD44 and PD-1 compared to WT T cells (Figure 1A, Extended Data Figure 1A). These differences translated to greater functionality as measured by increased cytokine production by PSGL-1^-/-^ T cells at 3 days post-activation (Extended Figure 1B). To directly and intrinsically address how TCR signaling was affected by PSGL-1 deficiency, we assessed the kinetics of Akt phosphorylation in WT and PSGL-1^*-/-*^ OT-I cells. TCR ligation was induced in a 1:1 mixture of WT and PSGL-1^-/-^ OT-I cells with cognate OVA_(257-264)_ peptide (SIINFEKL) and the cells were stimulated for the indicated times before fixation and analysis by phosflow flow cytometry (Figure 1B). We observed increased Akt activation early (15 mins) after TCR stimulation and sustained Akt phosphorylation (2 hr) in PSGL-1^-/-^ T cells.

**Figure 1.**
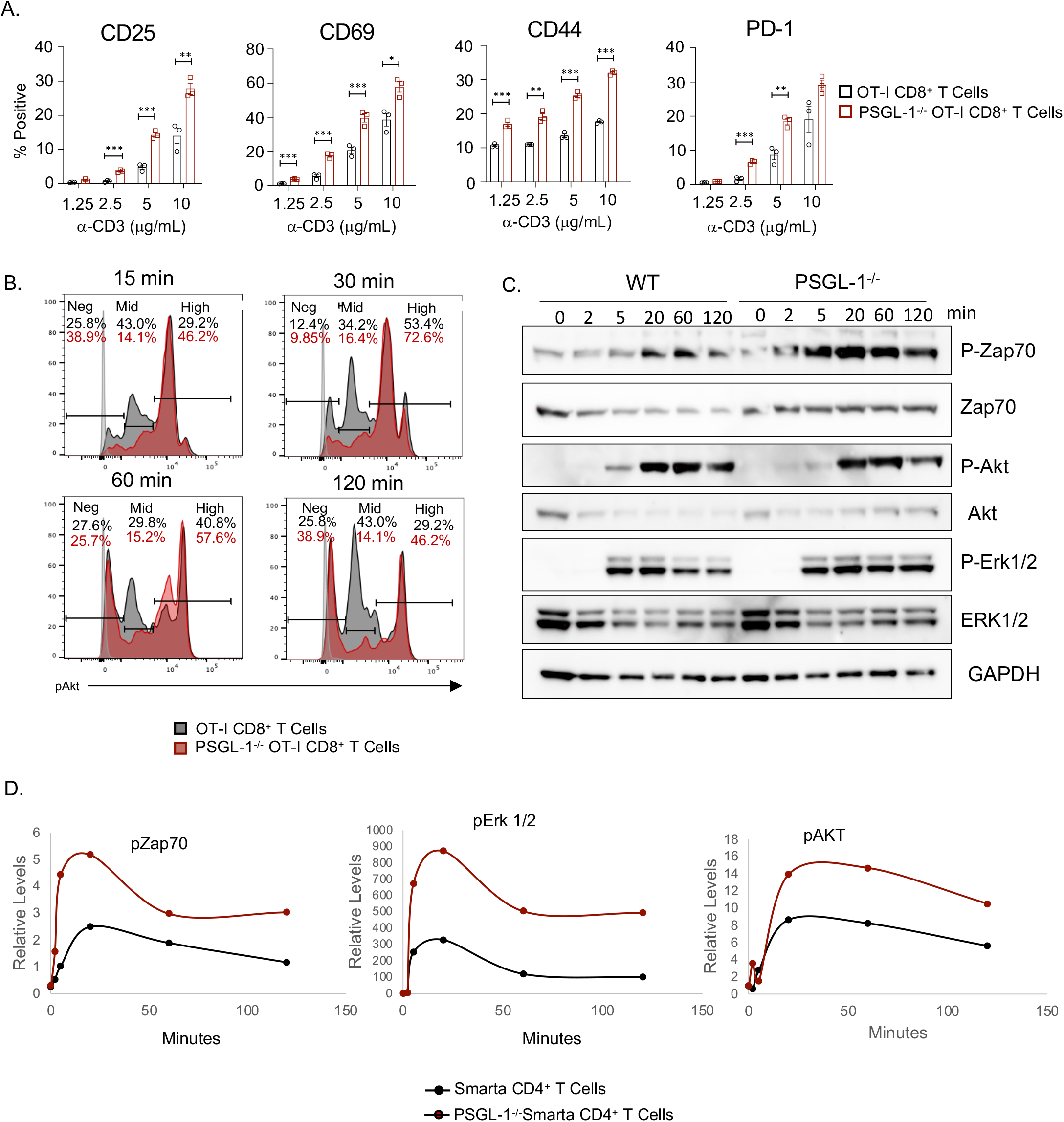
PSGL-1 restrains TCR stimulation strength and TCR signaling in T cells. (A) Bar graphs showing the frequency of CD8^+^ T cells expressing the activation markers: CD25, CD69, CD44^hi^, and PD-1 on OT-I WT and PSGL-1^-/-^ OT-I CD8^+^ T cells following overnight (18 hr) stimulation with plate-bound anti-CD3ε antibody at the indicated concentrations. Each dot represents an individual mouse. Data are representative of two independent experiments. p values are as follows: * < 0.05, ** < 0.01, *** < 0.005. Data are normally distributed. Unpaired t tests were used for statistical analysis of parametric data. (B) Representative flow cytometry histograms showing the kinetics of phosphorylated Akt expression in WT OT-I (dark gray) and PSGL-1^-/-^ OT-I (red) CD8^+^ T cells that were concomitantly stimulated with SIINFEKL OVA peptide-pulsed C57BL/6 splenocytes for the indicated times. Representative of 3 independent experiments. For each independent experiment, 2-3 mice per genotype were assessed. Light gray indicates FMO. (C) Western blot detection of phosphorylated and total levels of Zap70, Erk1/2, AKT and GAPDH in Smarta WT and PSGL-1^-/-^ CD4^+^ T cells stimulated for the indicated time with Streptavidin-conjugated anti-CD3ε antibody and crosslinking by anti-Strepavidin. (D) Line graphs showing relative levels of phosphorylated Zap70, Erk1/2, and AKT, relative to total protein expression for the cells analyzed in (C). Data are representative of one experiment from a pool of 2-3 mice per genotype.

To further address how PSGL-1 deficiency affected TCR signaling, we assessed proximal TCR signaling in WT and PSGL-1^*-/-*^ T cells by evaluating the phosphorylation kinetics of Zap70, Erk1/2 and Akt by Western blot (Figure 1C). Upon activation, PSGL-1^-^_/-_T cells demonstrated increased expression of pZap70, pErk1/2, and pAkt compared to WT T cells at each time point assessed and phosphorylation was sustained in PSGL-1^*-/-*^ T cells (Figure 1C, 1D). We also observed increased expression of these kinases with lower doses of anti-CD3 in PSGL-1^-/-^ T cells (Extended Data Figure 1C), demonstrating that PSGL-1 expression on T cells inherently limits T cells activation and the magnitude of T_EFF_ development from the time of initial activation.

Our previous study found that in exhausted CD8^+^ T cells, ligation of PSGL-1 simultaneously with TCR stimulation promoted greater phenotypic and functional exhaustion *in vitro* and *in vivo*^18^. To further test intrinsic regulation of TCR signaling by PSGL-1 under conditions of T cell exhaustion, we used an *in vitro* model of T cell exhaustion^23^ in which WT and PSGL-1^-/-^ OT-I T cells were repeatedly stimulated with SIINFEKL OVA peptide (*in vitro* T_EX,_ iT_EX_) and compared to effector CD8^+^ T cells that were optimally stimulated by a single peptide dose (*in vitro* T_EFF_, iT_EFF_). As we had observed that deficiency of PSGL-1 enables responsiveness to lower levels of TCR stimulation (Figures 1C-D, Extended Figure 1C), we hypothesized that PSGL-1 deficiency may also decrease TCR affinity sensitivity. We therefore used variant OVA peptides with single amino acid substitutions that result in differing affinities for the OT-I TCR to stimulate OT-I cells^24^. iT_EX_ and rested iT_EFF_ were assessed 5 days after initial activation with SIINFEKL peptide (N4), or with the lower affinity variants, SIIQFEKL (Q4, ∼18-fold decreased affinity), SIITFEKL (T4, ∼70-fold decreased affinity), or SIIVFEKL (V4, ∼700-fold decreased affinity). An equal frequency of WT and PSGL-1^-/-^ OT-I CD8^+^ T cells were activated (as determined by percent of cells with CD44 upregulation) by N4, Q4, and T4, while more PSGL-1^-/-^ OT-I CD8^+^ T cells were activated by V4 than WT OT-I CD8^+^ T cells (Figure 2A).

**Figure 2.**
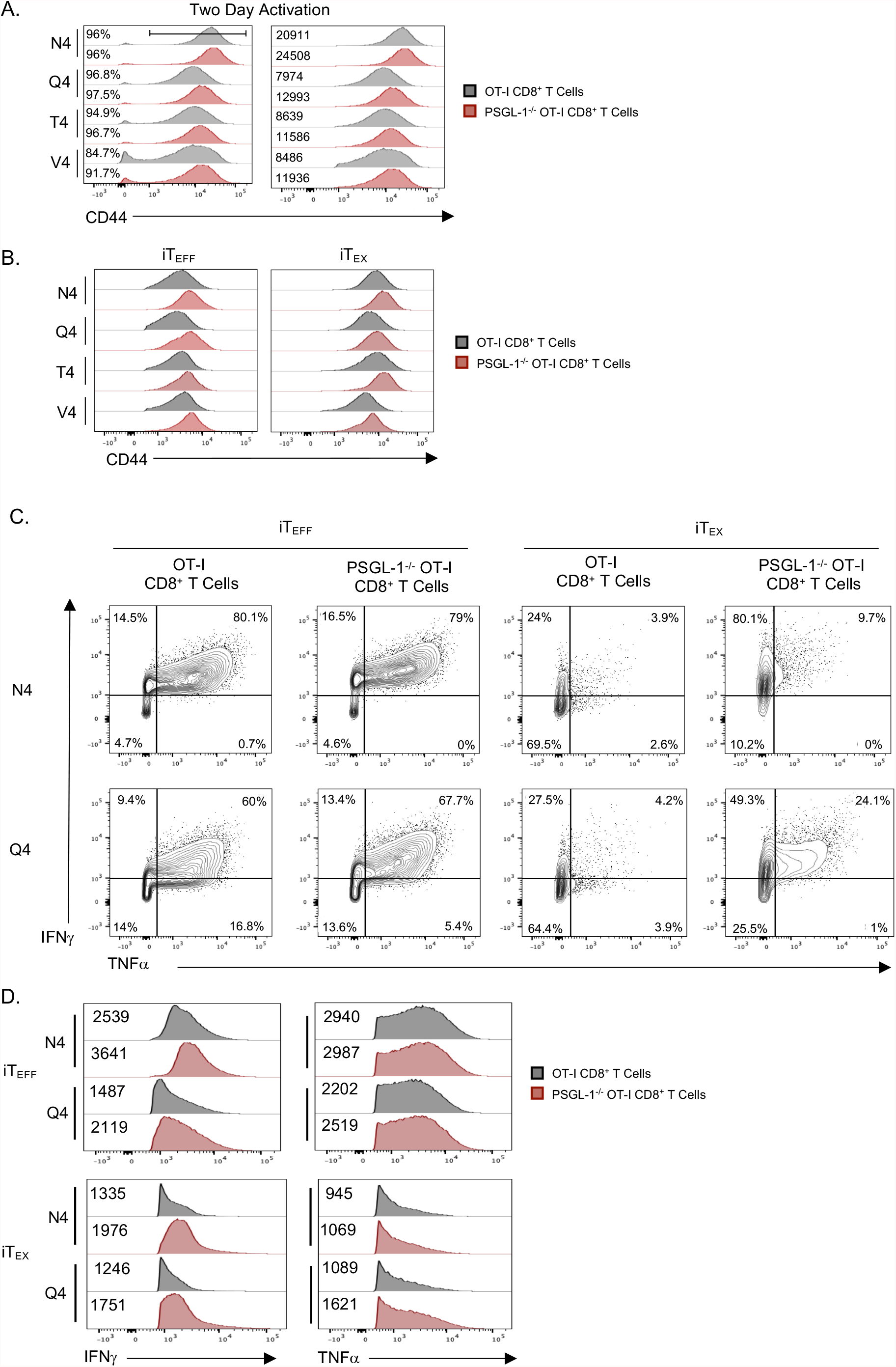
PSGL-1 deficiency promotes increased sensitivity to TCR signaling. OT-I and PSGL-1^-/-^ OT-I CD8^+^ T cells were cultured *in vitro* under effector conditions (single stimulation; iT_EFF_) or exhaustion conditions (repeated stimulation; iT_EX_) with SIINFEKL peptides of varying TCR affinity: N4 (SIINFEKL; regular affinity), Q4 (SIIQFEKL, ∼18-fold decreased affinity), T4 (SIITFEKL, ∼70-fold decreased affinity), or V4 (SIIVFEKL, ∼700-fold decreased affinity). (A) Left: Representative histograms showing the frequency of OT-I and PSGL-1^-/-^ OT-I CD8^+^ T cells that have upregulated CD44 by day 2 of activation with the indicated peptide in iT_EFF_ culture conditions. Right: Representative histograms showing expression levels of CD44 in OT-I and PSGL-1^-/-^ OT-I CD8^+^ T cells in iT_EFF_ on day 2 post-activation. (B) Representative histograms showing expression levels of CD44 in OT-I and PSGL-1^-/-^ OT-I CD8^+^ T cells in iT_EFF_ and iT_EX_ on day 5 post-activation. (C) Representative FACS plots of IFN^γ^ and TNFαproduction by OT-I and PSGL-1^-/-^ OT-I CD8^+^ T cells cultured with the indicated peptides under effector (left) or exhausted (right) conditions and restimulated on day 5 with SIINFEKL for 5 hours in the presence of Brefeldin A and monensin. (D) Representative histograms of IFNγ and TNFα production by OT-I and PSGL-1 ^-/-^ OT-I CD8^+^ T cells shown in (C). The numbers indicate the median fluorescence intensity (MFI) of each sample. Data in A-D are representative of three independent experiments. For each independent experiment, 2-3 mice per genotype were pooled.

At two days post-activation, despite a similar frequency of cells upregulating CD44 in response to N4, Q4 and T4 OVA stimulation, PSGL-1^-/-^ OT-I CD8^+^ T cells expressed higher levels of CD44 on a per cell basis (MFI, median fluorescence intensity); the same was true for V4-stimulated cells (Figure 2A). This pattern of increased CD44 expression was maintained in both iT_EX_ and rested iT_EFF_ conditions at day 5, when PSGL-1^-/-^ OT-I T cells exhibited greater activation than WT T cells stimulated with the original SIINFEKL peptide, N4, and with the progressively lower affinity peptides (Figure 2B). In order to assess the function of these cells, WT and PSGL-1^-/-^ OT-I iT_EFF_ and iT_EX_ cells generated with the N4 or Q4 peptides were restimulated with the high affinity N4 peptide and production of IFNγ and TNFα was evaluated by intracellular flow cytometry (Figure 2C). Restimulated WT *and* PSGL-1^*-/-*^ iT_EFF_ generated with either N4 or Q4 peptides had comparable frequencies of cells producing these cytokines, and most cells were double producers (WT: 59.70%; PSGL-1^-/-^: 57.37%)(Extended Figure 2A-B). However, PSGL-1^*-/-*^ iT_EFF_ demonstrated greater IFNγ production on a per cell basis (Figure 2D). WT T_EX_ generated from cultures with these peptides were highly exhausted, as evidenced by the very low frequencies of cytokine producers (N4: 2.6%, Q4: 3.9%)(Extended Data Figure 2A-B). PSGL-1^-/-^ T cells cultured under iT_EX_ conditions with the N4 peptide exhibited modest production of both IFNγ and TNFα (10.17%), Figure 2B upper right panel and Extended Data Figure 2A-B, and a 4.2-fold increase over WT iT_EX_ cultured with N4 peptide (Extended Data Figure 2A). In CD8^+^ T cells originally cultured with the lower affinity Q4 peptide, PSGL-1^*-/-*^ CD8^+^ T cells retained an increased capacity for double cytokine production (average 14.39%, Figure 2B lower right panel and Extended Data Figure 2A) with a 3.8-fold increase over WT iT_EX_ cultured with Q4 peptide (Extended Data Figure 2B). When evaluating overall cytokine production, the relative frequency of IFNγ-or TNFα-producing CD8^+^CD44^+^ T cells remained equivalent when comparing WT and PSGL-1^*-/-*^ cultured under iT_EFF_ conditions with either N4 or Q4 peptide (Extended Data Figure 2C-F). However, under iT_EX_ conditions, significantly more PSGL-1^*-/-*^ CD8^+^ T cells retained the ability to produce either IFNγ (N4: 2.3; Q4: 1.8) or TNFα (N4: 2.8; Q4: 2.1) relative to WT CD8^+^ T cells upon restimulation with N4 peptide (Extyended Data Figure 2C-F). Moreover, PSGL-1^*-/-*^ iT_EX_ showed greater IFNγ and TNFα production on a per cell basis (MFI)(Figure 2D). Taken together, these results show that PSGL-1 restrains T cell activation and early TCR signals required for iT_EFF_ development. Further, under conditions of repeated TCR stimulation leading to exhaustion, PSGL-1 limits responses to lower levels of TCR signal strength and lower affinity antigens, which would be detrimental to effective T cell responses, particularly in the context of lower affinity tumor cell antigens.

### Regulation of TCR signaling by PSGL-1

To more fully interrogate the regulation of TCR signaling by PSGL-1 more fully, we performed multiplexed mass spectrometry-based total and phospho-proteomics. Changes in the proteome and phosphoproteome of naïve WT and PSGL-1^-/-^ OT-I cells and OT-I cells stimulated for 2 hr with an optimal dose of immobilized anti-CD3 were assessed by mass spectrometry and the relative expression levels and significant differences of 7014 (total) and 9294 (phosphorylated) proteins were evaluated using MSstatsTMT. Upon activation, 576 phosphorylated proteins were significantly differentially expressed (≥ 2 log2FC, FDR < 0.01) between WT and PSGL-1^*-/-*^ OT-I CD8^+^ T cells (Extended Data Figure 3A). By employing ingenuity canonical pathway analysis comparing activated WT and PSGL-1^*-/-*^ OT-I CD8^+^ T cells, we identified 22 pathways in which significant (-log(p-value) ≥ 2) differences were predicted, including “Calcium-induced T Lymphocyte Apoptosis”, “CD28 Signaling in T Helper Cells”, “Nur77 Signaling in T Lymphocytes”, “iCOS-iCOSL Signaling in T Helper Cells”, “T cell receptor signaling”, and “PKCθ Signaling in T Lymphocytes” (Extended Data Figure 3B). We therefore evaluated changes in the phosphorylation state of proteins previously associated with regulation of T cell signaling/activation that were significantly (log2 fold change (p≤0.05)) upregulated in PSGL-1 deficient T cells (Extended Data Figure 3C), including Zap70, Lat, Stat5b, Myc, Nfatc2, Junb, and Fosl. Also notable were changes in the phosphorylation of proteins that regulate proliferation and survival (Extended Data Figure 3D) as well as calcium signaling, and metabolism (Extended Data Figure 3E). Gene set enrichment analysis confirmed our previous findings (Figure 1C-D) of the ERK1/2 pathway in PSGL-1^-/-^ OT-I cells (Extended Data Figure 3F). Together, these data support our conclusion that PSGL-1 signals play an essential role in the regulation of the T cell response to TCR engagement. Importantly, few changes were observed in total protein expression in naïve WT cells compared to naïve PSGL-1^-/-^ cells (30 up, 14 down; log2 fold change (p≤0.05)), with PSGL-1 being the third most differentially expressed protein (13.8-fold increase in WT cells). Of the 576 differentially expressed phosphoproteins, only 22% (127) were similarly differentially expressed in the naïve state.

Previous functional studies of PSGL-1 in T cell migration have indicated that this large, glycosylated protein engages ERM proteins along with actin filaments and migrates to the uropod that forms on the lagging side of the cells during polarized movement^25^. The localization of PSGL-1 with respect to the TCR signaling complex is therefore hypothesized to be outside the central area of TCR/peptide contact. To address the spatial relationship of PSGL-1 to the TCR complex, we analyzed their distribution by fluorescence microscopy. Naïve OT-I CD8^+^ T cells were incubated with either anti-CD3 or anti-PSGL-1 or both antibodies for 10 minutes at 37°C before being fixed and stained for PSGL-1 (red) and CD3 (green) expression localization (Figure 3A). In naïve cells, both PSGL-1 and CD3 were evenly distributed across the cell surface. Upon ligation with anti-CD3, CD3 formed punctate staining indicative of clustering as expected. Importantly, we observed that PSGL-1 co-clustered with CD3 under these conditions. When cells were ligated with anti-PSGL-1 antibody alone, PSGL-1 clustered to one side of the T cell while CD3 remained dispersed over the cell surface. However, when the cells were ligated simultaneously with both anti-CD3 and anti-PSGL-1, CD3 and PSGL-1 co-localized and migrated to one region of the cell. To quantify this co-expression in a high-throughput manner, we used imaging flow cytometry to evaluate expression and co-expression of CD3 and PSGL-1 on 10,000 CD8^+^ T cells per group treated in the same manner as above. The positive similarity score (Amnis software)(Figure 3B, Extended Data Figure 4A), confirmed and quantified the co-localization of CD3 and PSGL-1, supporting the hypothesis that PSGL-1 is optimally positioned to directly regulate the strength of TCR signals particularly when PSGL-1 is concomitantly engaged.

**Figure 3.**
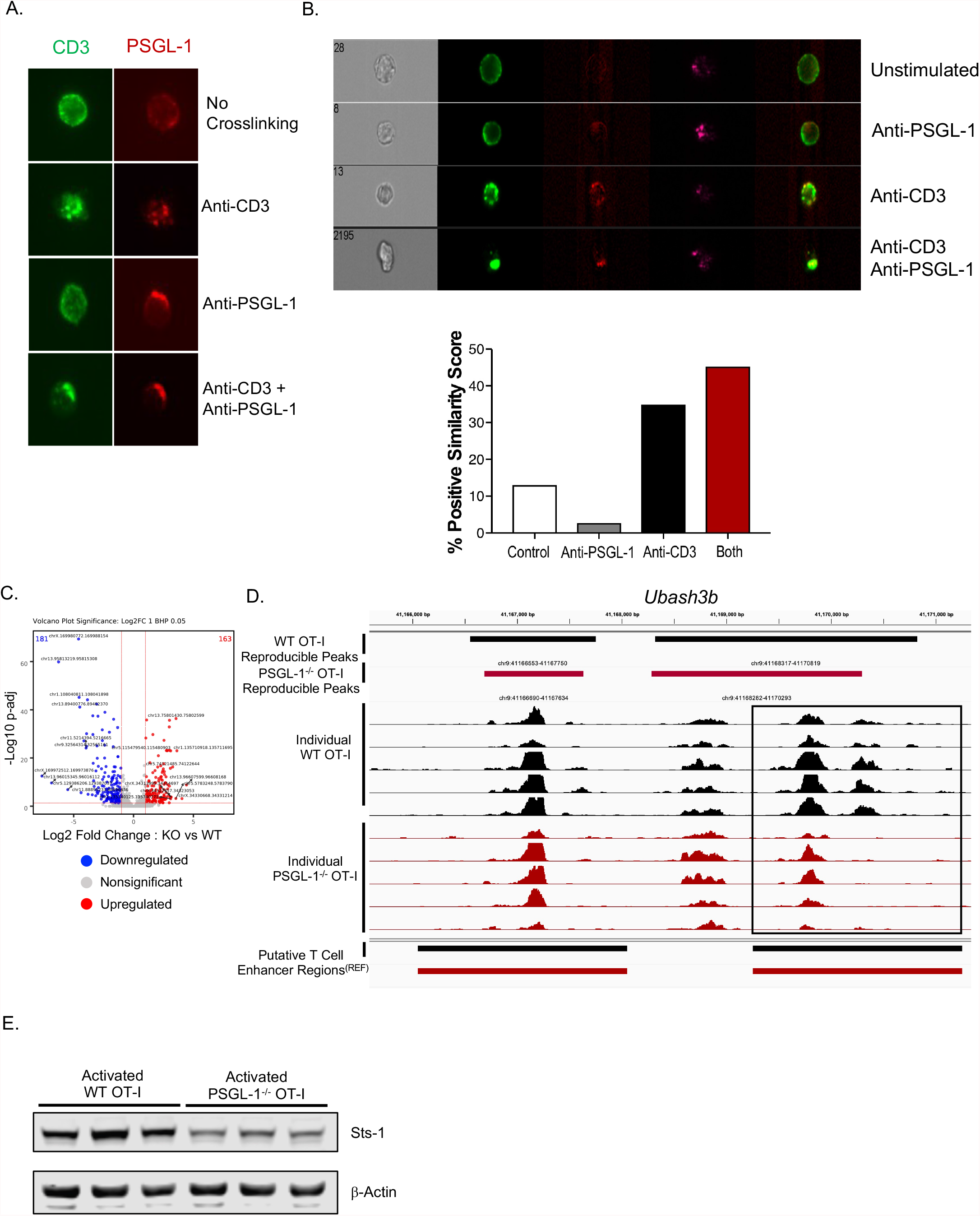
PSGL-1 co-localizes with the TCR and may promote TCR signaling attenuation via Sts-1. (A) Representative fluorescence microscopy images (40X) of single OT-I CD8^+^ T cells. Naïve OT-I CD8^+^ T cells were pre-stained with anti-CD3 and anti-PSGL-1 antibodies. Cells were then crosslinked as indicated to induce signaling in either CD3, PSGL-1, or both and incubated for 10 minutes at 37°C before fixation with (B) % formaldehyde. Localization was detected using SA-FITC (green, CD3) or goat anti-rat IgG (red, PSGL-1) and imaged using slides. Representative of two independent experiments. (B) Representative image of CD3 and PSGL-1 localization as described in A using an Amnis ImageStream Imaging Flow Cytometer. Imaging data from 10,000 individual cells was used to calculate the % positive similarity score. Representative of two independent experiments. Naïve OT-I CD8+ T cells were enriched from WT or PSGL-1^-/-^ splenocytes from 6-8 week old naïve OT-I mice followed by immobilized anti-CD3 (5 μg/mL, plate-bound) activation for two hours prior to the generation of ATAC-seq libraries. (C) Volcano plot of chromatin regions significantly (≥2 log2 FC, more open (upregulated) or closed (downregulated) in PSGL-1^-/-^ OT-I CD8^+^ T cells compared to OT-I CD8^+^ T cells. (D) Chromatin accessability within the *Ubash3b* gene region. WT OT-I tracks in black, PSGL-1^*-/-*^ OT-I tracks in red. Each individual track represents biological replicates prepared from two separate experiments. (E) Western blot analysis of Sts-1 and β-Actin loading control protein expression in two day activated WT OT-I and PSGL-1^-/-^ OT-I CD8^+^ T cells. Each band is an independent biologic replicate.

As we observed that PSGL-1^*-*/-^ CD8^+^ T cells were more readily activated and demonstrated increased expression of downstream TCR signaling molecules early after activation, we hypothesized that the epigenetic landscape would reflect this enhanced state of activation. Thus, we used ATAC-seq (Assay for Transposase Accessible Chromatin with high-throughput sequencing) to evaluate chromatin accessibility on a genome-wide level as an indication of changes in gene transcription that were regulated by PSGL-1. ATAC-seq was performed on WT and PSGL-1^-/-^ OT-I T cells after stimulation of naïve cells for 2 hr with an optimal dose of immobilized anti-CD3 (Extended Data Figure 4B-C). ATAC-seq analysis identified 163 chromatin regions that were significantly (Log2, p≤0.05) more open in PSGL-1^*-/-*^ OT-I T cells as compared to WT OT-I cells, and 181 chromatin regions that were more closed (Figure 3C). One gene identified was *Ubash3b*, which encodes the protein Sts-1 (Suppressor of T cell signaling-1), which was previously identified as a negative regulator of TCR signaling that inhibits activation of Zap70^20^. Evaluation of *Ubash3b* expression levels in CD8^+^ T cells in publicly available RNA-sequencing datasets (Immgen) showed that expression of *Ubash3b* increases in effector CD8^+^ T cells when compared to the naïve cells, suggesting that Sts-1 levels limit the extent of T cell activation as a means to control hyperresponsiveness of T cells^20^. However, what promotes or regulates Sts-1 expression in T cells has yet to be established. As part of a previous study, we performed mRNA RNA-sequencing on flow cytometry-based sorting of the immunodominant LCMV virus specific CD8^+^ T cells GP_(33-41)_ using MHC class I tetramers from LCMV Cl13 infected WT and PSGL-1^*-/-*^ mice^18^. We therefore evaluated whether *Ubash3b* was differentially expressed in GP_(33-41)_-specific CD8^+^ T cells in WT or PSGL-1^*-/-*^ mice. At 8 days post-infection with LCMV Cl13, RNA-sequencing analysis identified that *Ubash3b* is significantly downregulated (3.25 fold change, p ≤ 0.05) in PSGL-1^-/-^ T cells compared to WT T cells (GSE80113). ATAC-seq tracings identified an enhancer region within *Ubash3b* with significantly less open chromatin in PSGL-1^-/-^ T cells compared to WT T cells at 2 hr after activation with an optimal dose of anti-CD3 (Figure 3D). Western blot analysis further confirmed that naïve PSGL-1^*-/-*^ OT-I T cells express less Sts-1 (Extneded Data Figure 4D) and in activated CD8^+^ T cells (Figure 3E). Taken together, these data provide one potential mechanism for the enhanced TCR signaling observed within PSGL-1^-/-^ CD8^+^ T cells.

### PSGL-1 restrains glycolysis in CD8^+^ T cells

Given the effects of PSGL-1 on the extent of CD8^+^ T cell signaling and responses, we sought to identify its potential impact on glycolytic metabolism, which marks the switch from a resting, naïve state with low glycolysis to the highly glycolytic metabolic state that underlies T_EFF_ development and function upon TCR engagement^26^. Naïve WT or PSGL-1^-/-^ OT-I CD8^+^ T cells were activated for 3 days with immobilized anti-CD3 and anti-CD28 and glycolysis was assessed using the Seahorse Glycolytic Rate assay. The extracellular acidification rate (ECAR), a measure of glycolysis, and the oxygen consumption rate (OCR), a measure of mitochondrial oxidative phosphorylation, were assessed at baseline and upon the sequential addition of rotenone A and antimycin A, or 2-DG (Figure 4A). The proton efflux rate (PER), which measures proton efflux to extracellular media, and the glycolytic proton efflux rate (glycoPER) were also calculated (WAVE software, Agilent) (Figure 4B). At baseline, activated PSGL-1^-/-^ T cells exhibited significantly greater glycolytic activity compared to WT cells and remained at higher levels than WT cells upon the addition of rotenone A/antimycin A, indicating that this increased level of glycolysis was not simply due to increased mitochondrial activity despite overall greater oxygen consumption by PSGL-1^*-/-*^ OT-I T cells.

**Figure 4.**
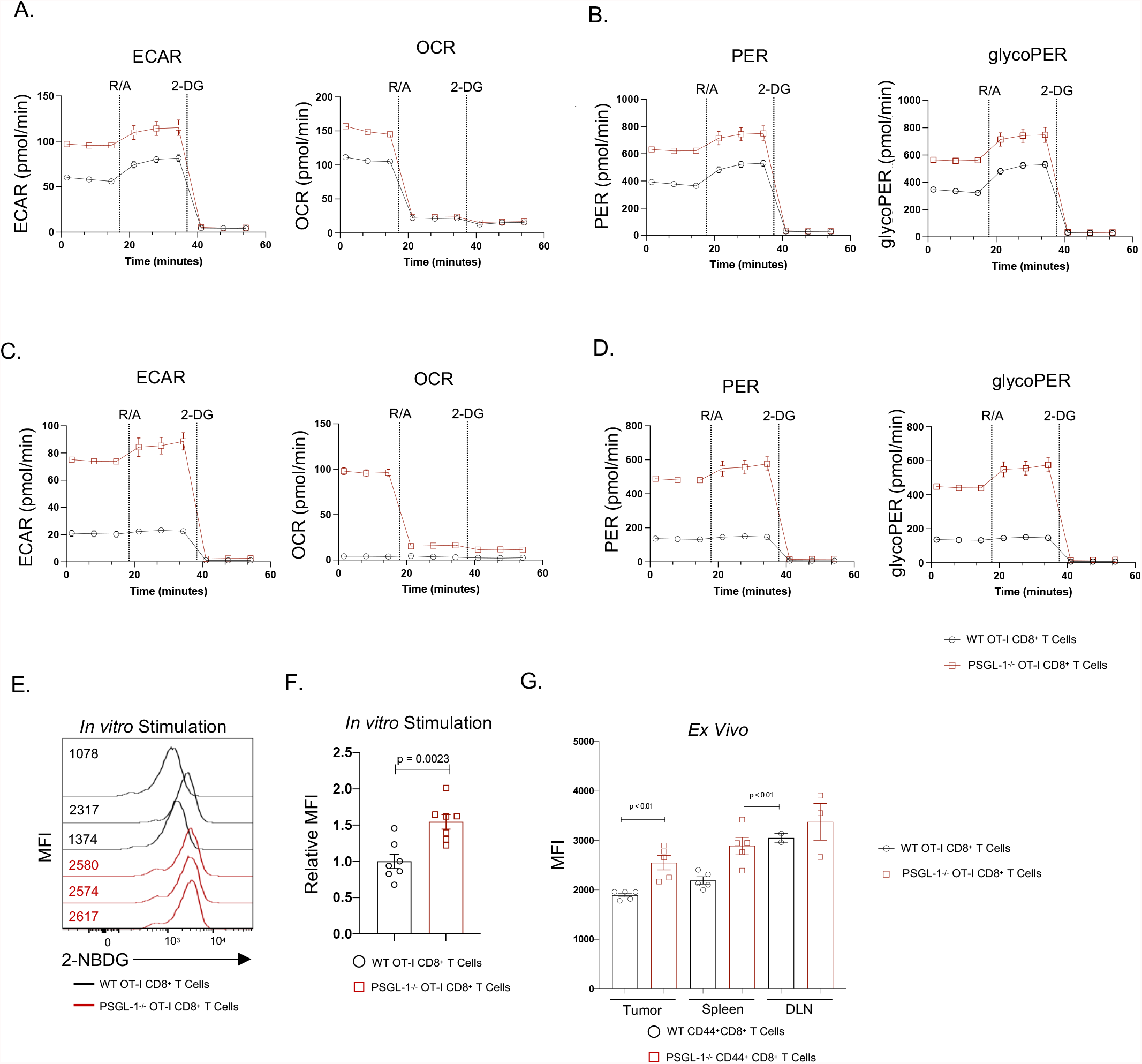
PSGL-1 restrains glycolysis in CD8^+^ T cells. Naïve WT or PSGL-1^-/-^ OT-I CD8^+^ T cells were activated for 3 days with plate-bound anti-CD3 and anti-CD28 (5 ug/mL, each) in the presence of 10 U/mL IL-2. (A) The extracellular acidification rate (ECAR) and oxygen consumption rate (OCR) were assessed using the Seahorse Glycolytic Rate Assay at baseline and following the sequential addition of rotenone A and antimycin A (R/A), or 2-deoxy-D-glucose (2-DG) as indicated. (B) The proton efflux rates (PER) and glycolytic rate (glycoPER) were calculated by WAVE software based on (A).and (B) representative of at least three individual experiments. *In vitro* exhausted OT-I or PSGL-1^-/-^ OT-I CD8^+^ T cells were generated as before with daily stimulation with SIINFEKL peptide 50 ng/mL in the presence of 5 ng/mL IL-7 and IL-15; glycolysis was assessed on day 5 using the Glycolytic Rate Assay. (C) ECAR and OCR were assessed using the Seahorse Glycolytic Rate assay at baseline and following the sequential addition of rotenone A and antimycin A, or 2-DG. (D) The proton efflux rates (PER) and glycolytic rate (glycoPER) were calculated by WAVE software based on (C). (C) and (D) are representative of at least three individual experiments. (E) Representative histogram of 2-NBDG uptake in OT-I or PSGL-1^-/-^ OT-I CD8^+^ T cells after 2 hr stimulation with SIINFEKL peptide; each line represents an individual mouse. (F) Dot plot/bar graph comparing relative 2-NBDG MFI in WT vs PSGL-1^-/-^ OT-I CD8^+^ T cells after 2 hr stimulation with SIINFEKL peptide. Each dot represents an individual mouse. Representative of three independent experiments. Data are normally distributed (Shapiro-Wilk test); an unpaired t test was used for analysis. (G) Dot plot/bar graph of *ex vivo* 2-NBDG MFI values in CD44^+^CD8^+^ T cells from tumors, spleens, or tumor draining lymph node (DLN) in WT or PSGL-1 mice bearing YUMM1.5 melanoma tumors. Data are normally distributed (Shapiro-Wilk) except for WT tumors and DLN. Unpaired t tests were used for statistical analysis of parametric data; Mann Whitney tests were used to statistical analysis of nonparametric data. Each dot in tumors and spleens represents an individual mouse; in DLN represents a pool of mice. Data are representative of one of two independent experiments.

Early during the development of T cell exhaustion, CD8^+^ T cells demonstrate reduced respiration (OCR) and glycolysis^27^. Further, inhibition of T_EX_ development such as through PD-1 blockade or genetic ablation is associated with an enhanced glycolytic phenotype and glucose uptake by CD8^+^ T cells^27^. As the *in vivo* phenotype of virus-specific PSGL-1^*-/-*^CD8^+^ T cells during chronic LCMV Cl13 infection is similar to PD-1 deficient (*Pdcd1*^-/-^) CD8^+^ T cells with increased expansion and functional capability, we hypothesized that PSGL-1^*-/-*^ CD8^+^T cells would similarly retain glycolytic activity under exhaustion conditions. We therefore sought to examine the glycolytic phenotype of exhausted PSGL-1^*-/-*^ T cells, we assessed glycolysis as above in *in vitro*-generated PSGL-1^*-/-*^ and WT exhausted OT-I CD8^+^ T cells (iT_EX_). As expected, phenotypically and functionally exhausted WT OT-I iT_EX_ cells have low levels of ECAR and OCR at baseline and upon mitochondrial uncoupling (Figure 4C). Comparatively, PSGL-1^*-/-*^ OT-I iT_EX_ retain high levels of ECAR and OCR (Figure 4C), corresponding to higher levels of PER and glycoPER than WT OT-I iT_EX_ (Figure 4D) and indicating that PSGL-1^-/-^ CD8^+^ T cells remain more glycolytically active under conditions of chronic TCR stimulation.

To address whether increased glycolytic metabolism of PSGL-1^*-/*-^ was accompanied by greater glucose utilization, we measured uptake of the fluorescent 2-NBD glucose analog (2-NBDG) by WT and PSGL-1^-/-^ OT-I T cells 2 hr after *in vitro* stimulation with SIINFEKL peptide by flow cytometry (Figure 4E). No differences in glucose uptake by naïve WT or PSGL-1^*-/-*^ OT-I T cells were observed, complementing our findings by Seahorse analysis that there were no differences in glycolysis in naïve CD8^+^ T cells (data not shown). We previously reported that PSGL-1 deficient mice exhibit significant control of the YUMM1.5 melanoma tumor line that is resistant to PD-1 blockade^18^. After initial tumor growth that was comparable to that in WT mice, we found that subsequent growth control was accompanied by much greater infiltration and effector function of CD8^+^ T cells^18^. To assess whether differences in glucose uptake could contribute to an enhanced antitumor response, we assessed 2-NBDG uptake by activated (CD44^hi^) T cells from YUMM1.5 tumor bearing mice. In CD44^hi^ CD8^+^ T cells from the tumors, spleens, and tumor draining lymph nodes (DLN) in WT or PSGL-1^*-/-*^ YUMM1.5 tumor-bearing mice assessed immediately *ex vivo*, we observed significantly greater capacity for glucose uptake by effector PSGL-1^*-/-*^ CD8^+^ T cells compared to WT effector CD8^+^ T cells (Figure 4G). Taken together, these data demonstrate that PSGL-1-deficient CD8^+^ TILs maintain a greater capacity for glucose uptake compared to WT CD8^+^ TILs and enhanced glycolytic capabilities under exhaustive conditions.

### scRNA-seq reveals distinct clustering of intratumoral PSGL-1 deficient CD8^+^ T cells and reveals an enhanced metabolic state

Previously we demonstrated that adoptively transferred PSGL-1^-/-^ OT-I T cells enabled tumor growth control of B16-OVA in C57BL/6 recipients to a significantly greater extent than WT OT-I cells^18^. To further assess intrinsic differences between WT and PSGL-1^-/-^ T cells *in vivo*, we used single cell RNA-seq (scRNA-seq) and the B16-OVA melanoma model to identify changes at the transcriptional level. Seven days after B16-OVA tumor cell inoculation into C57BL/6 mice, *in vitro* activated WT or PSGL-1^*-/-*^ OT-I CD8^+^ T cells were adoptively transferred into tumor-bearing mice. Donor OT-I cells were then sorted from the tumor-draining inguinal lymph node or B16-OVA tumors 3 or 6 days post-transfer, respectively and (scRNA-seq) libraries were generated using the 10X Genomics platform (Figure 5A). CellLoupe (10X Genomics) t-distributed stochastic neighbor embedding (t-SNE) analysis shows distinct clustering of WT and PSGL-1^-/-^ OT-I TILs (Figure 5B). Seurat-based tSNE analysis displayed a greater mix of WT and PSGL-1^*-/-*^ OT-I TILs across all clusters, yet a heatmap (DoHeatmap, Seurat) displaying the top most differentially expressed genes (based on average log2FC) identified pockets of genes that were globally differentially expressed in PSGL-1 deficient cells (*Tmsb10, Crip1, Ifi27l2a, Mt1, Mt2, Atp5k*) as well as cluster-specific alterations in genes (Figure 5C). To further interrogate differential gene expression in effector CD8^+^ T cells, we used SeqGeq (Flow Jo, BD) to identify OT-I cells co-expressing granzyme B (*Gzmb*) and IFNγ (*Ifng*). We observed that more PSGL-1^-/-^ OT-I cells were co-expressing *Gzmb* and *Ifng and* this co-expression was linked to greater expression of *Mtor* and *Hif1a* as well as engagement in cell cycle (*Mki67*), but without changes in the regulation of cell survival as measured by *Bcl2* (Figure 5D).

**Figure 5.**
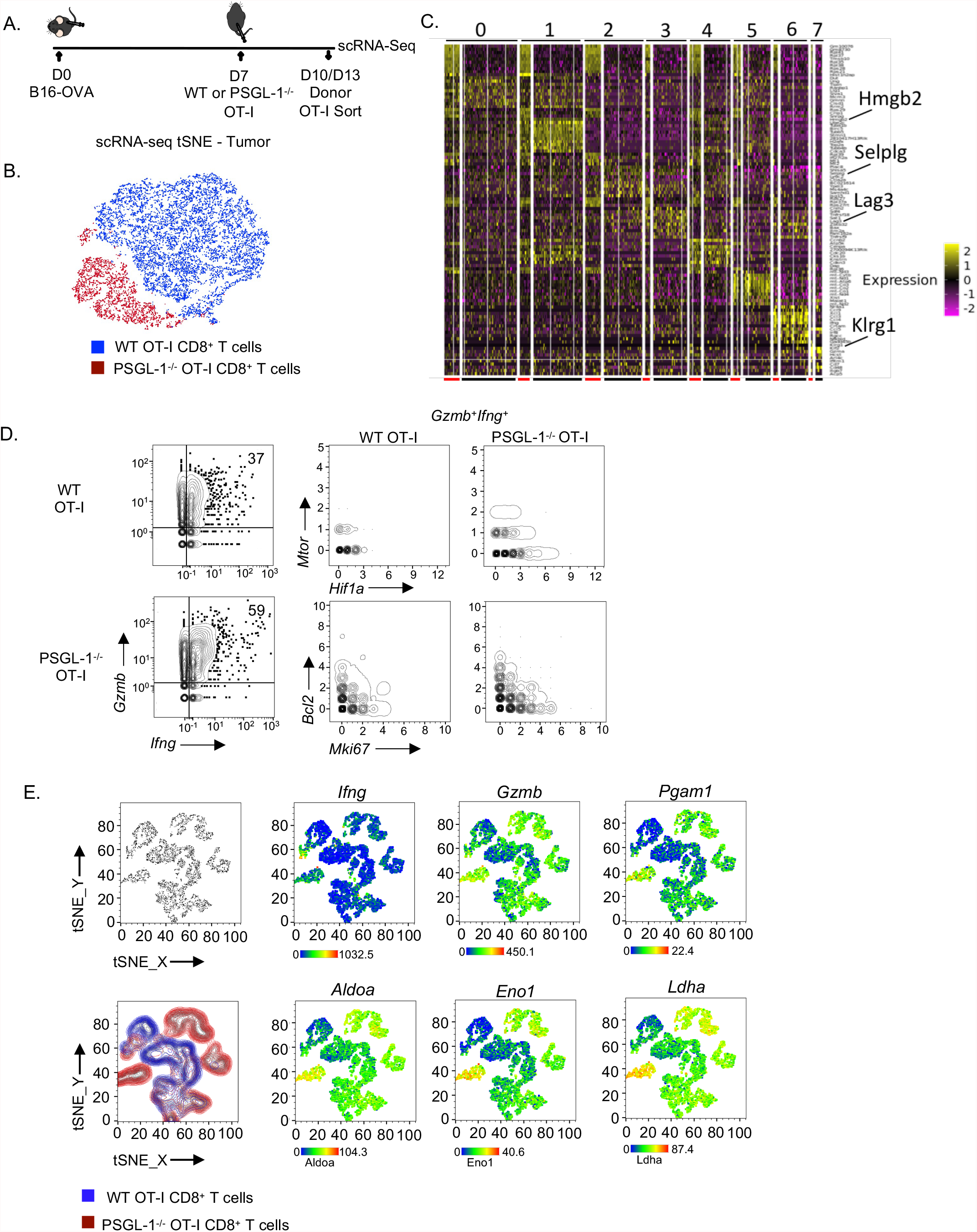
Single-cell sequencing reveals enhanced metabolic state of intratumoral CD8+ T cells in the absence of PSGL-1. (A) Experimental design for the generation of scRNA-seq libraries. C57BL/6 (CD45.2/CD90.2) mice received a subcutaneous injection of 1×10^6^ B16-OVA melanoma tumor cells. Tumors were allowed to become established and on day 7 mice received either 1×10^6^ *in vitro* activated OT-I (CD45.1^+^) or PSGL-1^*-/-*^ OT-I (CD90.1^+^) CD8+ T cells. Three days after T cell injection (day 10 overall), donor OT-I cells were FACS sorted from the inguinal tumor draining and non-draining lymph nodes. Six days after T cell injection (day 13 overall), donor OT-I cells were FACS sorted from the inguinal tumor draining lymph nodes or B16-OVA tumors from a separate subset of mice. 10X Genomics single-cell RNA-sequencing (scRNA-seq) libraries were prepared immediately following FACS sorting. (B) 10X CellLoupe-defined tSNE cluster of OT-I and PSGL-1^*-/-*^ OT-I CD8^+^ T cells sorted from B16-OVA tumors. (C) Seurat-generated heatmap showing the top 100 most significant differentially expressed genes between WT and PSGL-1^-/-^ OT-I CD8^+^ T cells as determined by scRNA-seq. Each row reflects an individual gene and each column an individual cell. The top numbers indicate the corresponding cluster of each cell and the red (PSGL-1^-/-^) and black (WT) lines indicate the genotype of the cell. (D) SeqGeq analysis of *Gzmb* and *Ifng* expression in WT and PSGL-1^*-/-*^ OT-I CD8^+^ T cells (left). Top right: gene expression of *Mtor* and *Hif1a* in *Gzmb*^*+*^*Ifng*^*+*^ WT and PSGL-1^*-/-*^ OT-I CD8^+^ T cells. Bottom right: gene expression of *Bcl2* and *Mki67* in *Gzmb*^*+*^*Ifng*^*+*^ WT and PSGL-1^*-/-*^ OT-I CD8^+^ T cells. (E) SeqGeq tSNE clustering of WT and PSGL-1^*-/-*^ OT-I CD8^+^ T cells (top) with library source color overall (bottom); WT, blue; PSGL-1^*-/-*^, red. Gene expression overlays of *Ifng, Gzmb, Pgam1, Aldoa, Eno1*, and *Ldha*. scRNA-seq data representative of over 1000 single cells per condition pooled from 6 mice per group.

As we identified greater capacity for glycolysis with PSGL-1 deficiency by OT-I cells, we assessed expression of genes associated with glycolysis in WT (blue) and PSGL-1^-/-^ (red) OT-I cells by overlaying gene expression on a SeqGeq-generated tSNE plot (Figure 5E), which allows the “gating” and evaluation of gene expression within a subset of scRNA-seq gene expression data. Individual *Ifng* and *Gzmb* expression was largely localized in PSGL-1^*-/-*^ OT-I T cell clusters. Furthermore, the greatest expression of the genes *Pgam1, Aldoa, Eno1*, and *Ldha* was co-localized with these cytotoxic-capable PSGL-1^*-/-*^ OT-I T cells. These analyses underscore that PSGL-1 deficiency confers greater metabolic capacity in terms of glycolysis even in the context of the tumor microenvironment. Further, the results demonstrate that greater metabolic capacity is linked to greater effector function, identifying that the absence of PSGL-1 releases metabolic constraints that are essential for efficient CD8^+^ T cell effector functions, particularly in the tumor microenvironment.

### PSGL-1 suppresses the generation of stem cell-like T_EX_ and promotes TOX expression

Studies using the LCMV model of chronic infection have identified stem-cell-like T_EX_ (designated T_SC_) with a capacity for self-renewal as well as capacity for expansion and effector function^28-30^. These CD8^+^ T cells are defined by high expression of the transcription factor TCF-1. In metastatic melanoma, TCF-1^+^ CD8^+^ T cells have been identified in patient tumors with objective responses to PD-1 blockade^21^. Several reports have now shown that the transcription factor, TOX, is a fundamental regulator that ultimately determines a T cell’s fate of irreversible exhaustion^31-34^. However, co-expression of TOX and TCF-1 has been shown to distinguish less exhausted T cells that can retain effector function^32,34^. Since PSGL-1 deficiency was associated with greater T cell function in our studies of LCMV Cl13, we assessed TCF-1 and TOX expression in WT and PSGL-1^-/-^ GP_(33-41)_^+^ virus-specific CD8 ^+^T cells by tetramer staining in the spleens at 15 days after infection with LCMV Cl13 (Figure 6A, B) when the virus is cleared from the sera of PSGL-1^-/-^ GP_(33-41)_^+^ mice^18^. A significantly higher frequency of TCF-1^+^ cells was found in PSGL-1^-/-^ mice compared to WT mice, and conversely, a large fraction of WT GP _(33-41)_^+^ T cells expressed TOX, compared to a small percentage of PSGL-1^-/-^ GP_(33-41)_^+^ T cells. We then addressed whether differences in expression of these transcription factors were present in GP _(33-41)_^+^ and NP _(396-404)_^+^ virus-specific cells at 9 days after infection of WT and PSGL-1^-/-^ mice when the virus loads are comparable, as compared to 15 days when only WT mice display virus in the blood. Among GP _(33-41)_^+^ T cells from the blood of LCMV Cl13 infected animals, the frequencies of TCF-1^+^ cells were indistinguishable at 9 dpi between WT and PSGL-1^*-/-*^ mice but dramatically fewer GP _(33-41)_^+^ T cells expressed TOX (Figure 6C). At day 15, the phenotype of GP_(33-41)_^+^ T cells from the blood reflected those in the spleen, with a moderate increase in TCF-1^+^ cells and a drastic decrease in TOX^+^ cells (Figure 6C). In both WT and PSGL-1^-/-^ LCMV Cl13-infected mice, the majority of GP_(33-41)_^+^ T cells were both TCF-1^-^ and TOX^-^ (Figure 6C and Extended Data Figure 5A). Evaluation of IL-7Rα and KLRG-1 expression on CD8^+^ T cells classically defines the delineation between short-lived effector cells (SLEC) and memory precursor cells (MPEC) in response to some infections, including LCMV Arm infection^35^, and we previously observed during LCMV Arm infection that GP_(33-41)_-specific PSGL-1^-/-^ CD8^+^ T cells skew towards the MPEC phenotype^19^. Previous studies have demonstrated an inverse relationship between TOX and KLRG1 expression during LCMV Cl13 virus infection^36^. In our study, we also observed that all KLRG1^+^ cells were within the TOX^-^ population for both WT and PSGL-1^-/-^ GP_(33-41)_-specific CD8^+^ T cells. When we evaluated the differentiation of TOX^-^ GP_(33-41)_-specific CD8^+^ T cells, we found that there were fewer early effector T cells (IL-7Rα^−^/KLRG-1^-^) PSGL-1^-/-^ CD8^+^ T cells, and increased representation within the IL-7Rα^+^/KLRG-1^-^ and IL-7Rα^+^/KLRG-1^+^ populations (Extended Data Figure5A-B).

**Figure 6.**
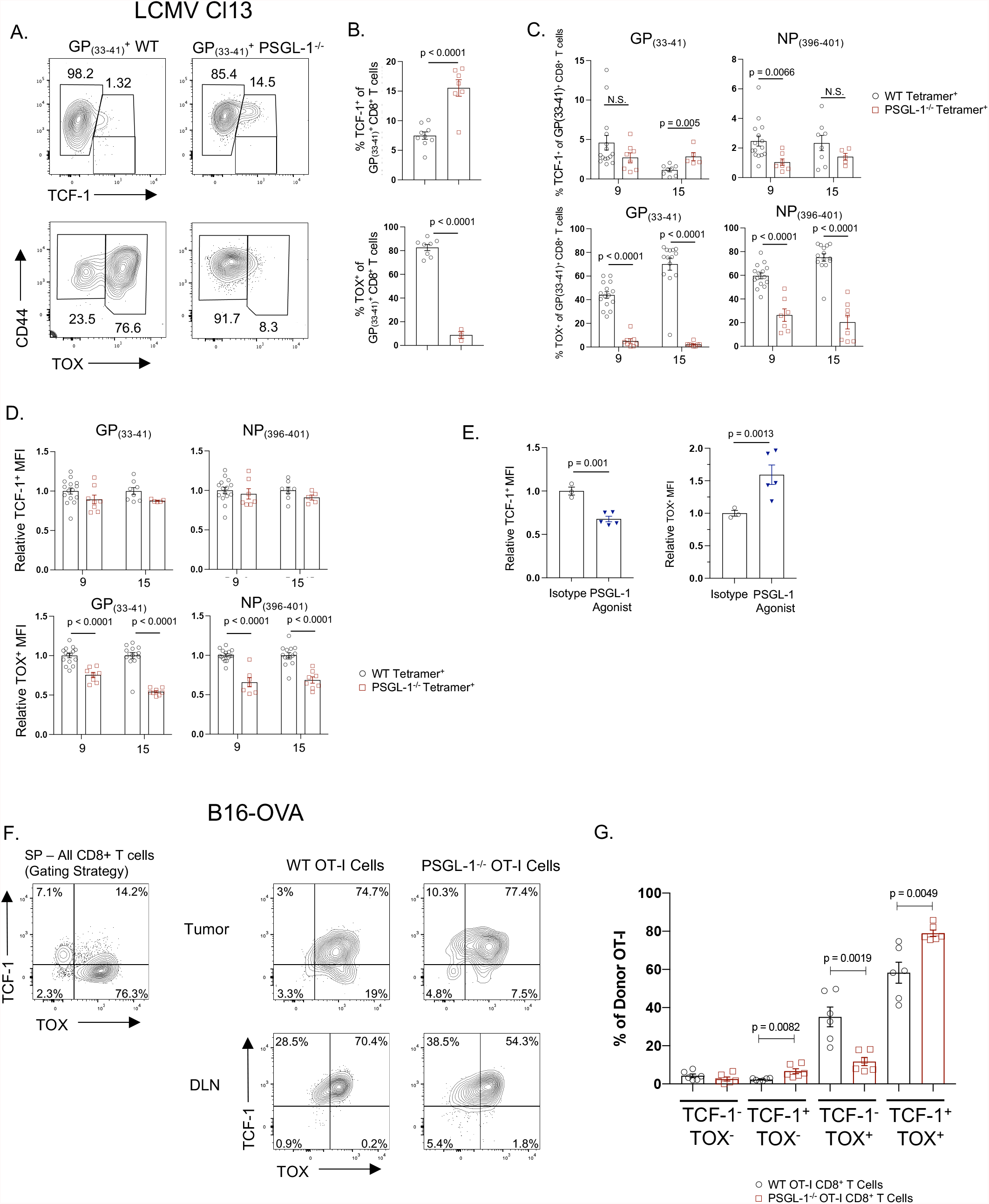
PSGL-1 limits TCF-1 and promotes TOX expression in CD8^+^ T cells during chronic virus infection and cancer. C57BL/6 WT or PSGL-1^-/-^ mice were infected with LCMV Cl13 virus. Virus-specific T cell responses were assessed using GP_(33-41)_^+^ and NP_(396-404)_^+^MHC-I tetramers in combination with flow cytometry. (A) Representative flow cytometry plots showing TCF-1 or TOX expression vs CD44 expression in GP_(33-41)_- specific CD8^+^ T cells from the spleens of WT or PSGL-1^*-/-*^ mice on day 15 post-infection. (C) Dot plot/bar graphs of the frequency of TCF-1^+^ (top) or TOX^+^ (bottom) GP_(33-41)_^+^ CD8 ^+^ T cells in WT or PSGL-1^*-/-*^ mice on day 15 post-infection. Data are normally distributed (Shapiro-Wilk); unpaired t test was used for statistical analysis. TCF-1 data are from two independent experiments, TOX data are from one experiment. Each dot represents an individual mouse. (C) Dot plot/bar graphs of the frequency of TCF-1^+^ (top) or TOX^+^ (bottom) GP_(33-41)_^+^ or NP_(396-404)_^+^ CD8^+^ T cells in the blood of WT or PSGL-1^*-/-*^ mice on days 9 and 15 post-infection. Data are normally distributed (Shapiro-Wilk) except for D9 WT GP_(33-41)_ and NP_(396-404)_ TCF-1, D15 WT GP_(33-41)_ and NP_(396-404)_ TOX, D9 and D15 KO GP_(33-41)_ TOX. Unpaired t tests were used for statistical analysis of parametric data; Mann Whitney tests were used to statistical analysis of nonparametric data. Data in C are from at least three independent experiments. Each dot represents an individual mouse. (D) Dot plot/bar graphs of the relative per-cell expression (MFI) of TCF-1^+^ (top) or TOX^+^ (bottom) GP_(33-41)_ or NP_(396-404)_ CD8 T cells in the blood of WT or PSGL-1 ^*-/-*^ mice on days 9 and 15 post-infection. Data are normally distributed (Shapiro-Wilk) except for D15 WT GP_(33-41)_ TOX, and D9 KO NP_(396-404)_ TCF-1. Unpaired t tests were used for statistical analysis of parametric data; Mann Whitney tests were used to statistical analysis of nonparametric data. Data in (D) are from at least three independent experiments. Each dot represents an individual mouse. WT C57BL/6 mice were infected with LCMV Cl13, and were treated with either isotype control antibody or a PSGL-1 agonist antibody (4RA10) on days 5 and 8 post-infection. (E) Dot plot/bar graphs of the relative per-cell expression (MFI) of TCF-1^+^ (top) or TOX^+^ (bottom) GP_(33-41)_^+^ or NP _(396-404)_^+^ CD8^+^ T cells in the spleens of isotype or PSGL-1 agonist-treated mice on day 15 post-infection. Data are normally distributed (Shapiro-Wilk). Unpaired t tests were used for statistical analysis of parametric data. Data in E are representative of one of two independent experiments. Each dot represents an individual mouse. WT C57BL/6 (CD45.2/CD90.2) mice were inoculated subcutaneously with 1×10^6^ B16-OVA melanoma tumor cells. Two weeks later, mice received either 1×10^6^ *in vitro* activated WT OT-I (CD45.1^+^) or PSGL-1^*-/-*^ OT-I (CD90.1^+^) CD8^+^ T cells. 6 days later, donor OT-I cells in B16-OVA tumors or inguinal tumor draining lymph nodes (DLN). (F) TCF-1 and TOX expression in CD44^hi^ donor OT-I CD8^+^ T cells was assessed by intracellular flow cytometry. (G) Dot plot/bar graphs of TCF-1 and TOX expression in donor OT-I CD8^+^ T cells in the tumors of B16-OVA tumor-bearing mice. Data are normally distributed (Shapiro-Wilk). Unpaired t tests were used for statistical analysis of parametric data. Data in G are representative of one of two independent experiments. Each dot represents an individual mouse.

Unlike GP_(33-41)_-specific CD8^+^ T cells which persist throughout LCMV Cl13 infection, CD8^+^ T cells recognizing the higher avidity immunodominant LCMV epitope NP_(396-404)_ are commonly deleted over the course of chronic LCMV Cl13 infection^37,38^, but are retained at a higher frequency in PSGL-1^-/-^ mice^18^. In the high affinity NP _(396-404)_^+^ T cells, fewer PSGL-1^-/-^ T cells expressed TCF-1 than WT cells at day 9 with no significant difference by day 15, but a lower frequency of PSGL-1^-/-^ T cells expressed TOX at both day 9 and day 15 post-infection (Figure 6C). Further, while expression levels of TCF-1 on a per cell basis (MFI) were indistinguishable for either GP_(33-41)_^+^ or NP_(396-401)_^+^ T cells, TOX expression levels were consistently lower in PSGL-1^-/-^ virus-specific T cells at both day 9 and day 15 (Figure 6D). Together these data support the conclusion that PSGL-1 deficiency prevents terminal differentiation of T_EX_, in part by limiting the expression of TOX.

To test the hypothesis that PSGL-1 engagement is a driver of TOX expression and T_EX_ differentiation, we assessed TCF-1 and TOX expression *ex vivo* in virus-specific GP _(33-41)_^+^ CD8^+^ T cells from LCMV Cl13-infected WT mice treated with either an agonist anti-PSGL-1 antibody (clone 4RA10) which drives T cell exhaustion^18^ or an isotype control antibody. PSGL-1 ligation in the context of LCMV Cl13 infection reduced the per-cell expression of TCF-1 by 25%, while TOX expression was increased by 50% following PSGL-1 ligation during infection (Figure 6E). To address T cell intrinsic effects of PSGL-1 regulation on TCF-1 and TOX expression in response to melanoma tumors, we analyzed *in vitro* activated and adoptively transferred WT and PSGL-1^-/-^ OT-I cells 10 days after transfer into B16-OVA tumor-bearing WT mice. We observed a modest increase in the frequencies of TCF-1^+^CD44^+^OT-I^+^ T cells in B16-OVA tumors from PSGL-1^*-*/-^ mice (Figure 6F, top right panel) compared to the tumors from WT mice (Figure 6F, top left panel). In the inguinal tumor-draining lymph node, essentially all donor OT-I cells were TCF-1^+^, but fewer cells co-expressed TOX in PSGL-1^*-/-*^ donor OT-I CD8^+^ T cells. Evaluation of single and co-expression of TCF-1 and TOX in donor OT-I CD8^+^ T cells in B16-OVA tumors found that very few donor OT-I cells lack expression of both TCF-1 and TOX. PSGL-1^*-/-*^ OT-I CD8^+^ TILs displayed a slight increase in TCF-1^+^ single-positive (SP) cells, significantly decreased TOX^+^ SP cells and increased TCF-1/TOX double-positive (DP) cells compared to WT donor OT-I CD8^+^ T cells (Figure 6G). These data suggest that a substantial proportion of tumor-specific TEX cells were progressively differentiating during the response within the tumors. In addition, decreased expression of TOX consistently distinguished PSGL-1^-/-^ T cell in the settings of chronic antigen stimulation, confirming a fundamental role for PSGL-1 in promoting T cell exhaustion.

### Modulation of PSGL-1 signaling can promote or block T cell exhaustion

Given our findings in murine models of chronic infection and cancer wherein *in vivo* PSGL-1 ligation promotes CD8^+^ T cell exhaustion during LCMV Cl13 infection^18^ and increased tumor growth (data not shown), we next sought to address whether PSGL-1 can similarly regulate the function of human CD8^+^ T cells. We therefore established an *in vitro* model of human T_EFF_ and T_EX_ differentiation from healthy donor PBMCs using single (iT_EFF_) or repeated (iT_EX_) TCR stimulation with tetrameric anti-CD3/CD28 complexes (ImmunoCult, StemCell) which generates polyfunctional IFNγ- and TNFα-producing iT_EFF_ and iT_EX_ that fail to produce cytokines upon restimulation with anti-CD3/CD28 complexes. Restimulation of T cells 9 days after culturing under iT_EFF_ conditions resulted in CD45RO^+^CD8^+^ T cells with a robust capacity to individually produce IFNγ and TNFα, as well as to co-produce these cytokines (average 17.3%)(Figure 7A left panels and Figure 7B). In contrast, restimulation of T cells cultured under iT_EX_ conditions failed to produce significant amounts of either cytokine (average 0.8%)(Figure 7A right panels and Figure 7B). Previous studies have linked the relative expression of the transcription factors T-box expressed in T cells (T-bet) and Eomesodermin (Eomes) to T cell exhaustion such that terminally exhausted iT_EX_ express lower levels of T-bet and increased levels of Eomes compared to iT_EFF_ in chronic virus infection in mice with LCMV Cl13^39^, in HIV infected patients^40^, and in the context of cancer^41^. In our *in vitro* model, we found that the relative Eomes/T-bet expression (based on relative MFI) increased in iT_EX_ cells compared to iT_EFF_ (Figure 7C). When combined with the decreased functional capacity of these iT_EX_ cells, this indicates that these *in vitro* cultured cells have a phenotype reflective of T_EX_ from patients. This is particularly relevant for human T cells, as TOX expression is not limited to T_EX^42^_. To address how PSGL-1 signaling in human T cells may contribute to CD8^+^ T cell differentiation, iT_EX_ cells were additionally cultured in the presence of a PSGL-1 agonist antibody (KPL-1). In 4 out of 5 donors assessed, one time activation of T cells in the presence of a PSGL-1 agonist antibody resulted in the decreased ability of these cells to produce and co-produce IFNγ and TNFα upon restimulation 9 days after initial activation (Figure 7A middle panels and Figure 7B). For those donors in which cytokine production dropped, capacity for co-production was an average 55% of the restimulated effector cells (Figure 7B). Consistent with an incomplete differentiation of iT_EX_ from iT_EFF_ when PSGL-1 ligation was present together with single TCR stimulation, we observed a modest increase in the relative Eomes/T-bet expression ratio (relative MFI)(Figure 7C). When combined with repeated TCR stimulation, the Eomes/T-bet expression ratio of iT_EX-_treated CD8^+^ T cells increased to a similar level as iT_EFF_ without PSGL-1 ligation (Figure 7C). These data indicate that signaling via PSGL-1 concomitant with TCR stimulation can intrinsically orchestrate the development of T_EX_ from T_EFF_ in human CD8^+^ T cells.

**Figure 7.**
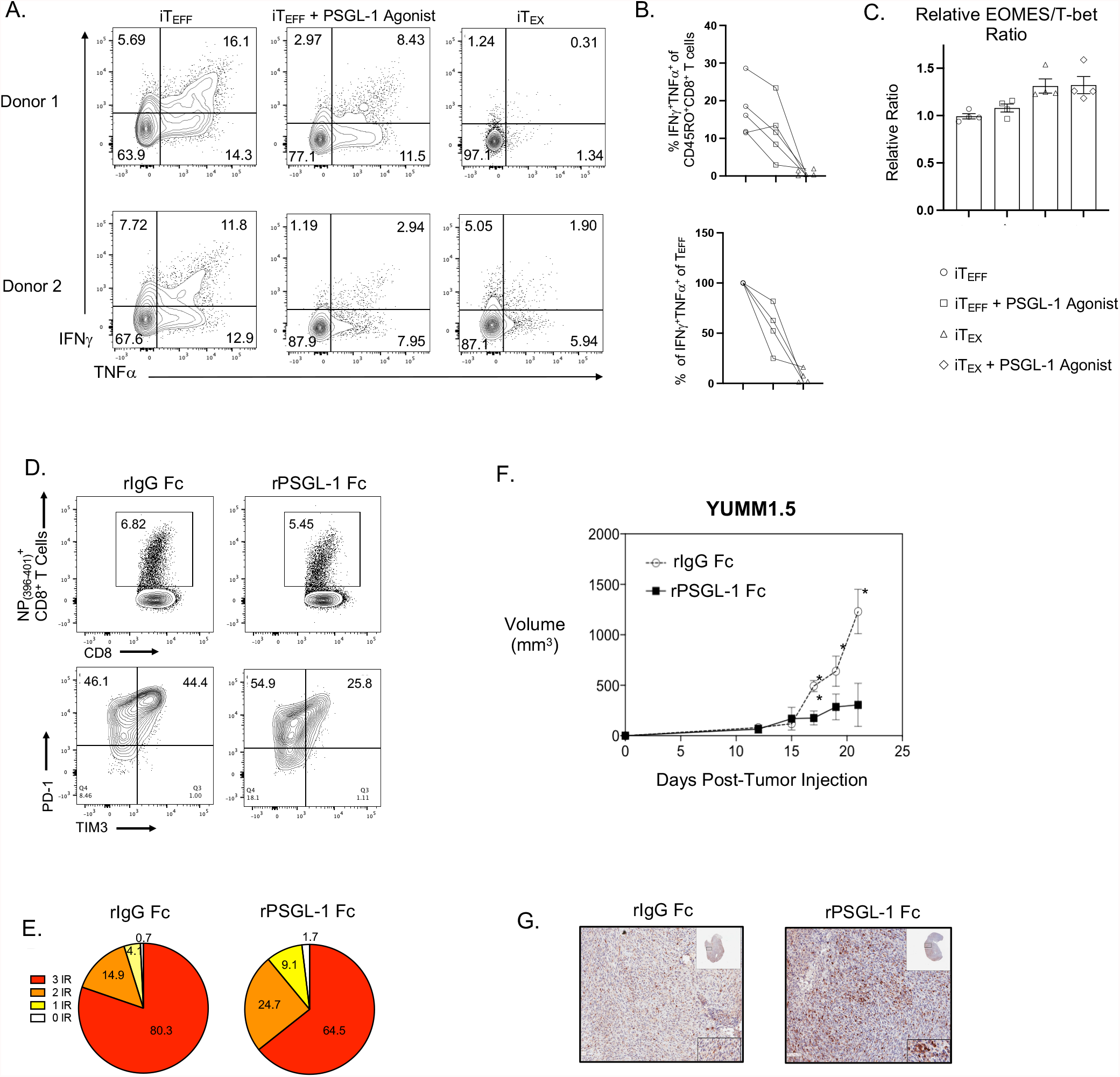
Pharmacological inhibition of PSGL-1 promotes decreased T cell exhaustion and functional T cell responses to melanoma tumors and LCMV Cl13. PBMCs from healthy donors were cultured under effector (single stimulation, iT_EFF_) or exhausted (repeated stimulation, iT_EX_) conditions for 9 days and then assessed immediately for transcription factor expression or restimulated overnight to assess cytokine production by intracellular flow cytometry. (A) Representative flow cytometry plots showing IFNγ and TNFαproduction by PBMCs from two different donors following restimulation. Plots are pre-gated on live, CD8^+^CD45RO^+^ cells. (B) Top: Dot plots showing the frequency of IFNγ and TNFα double-producing CD8^+^CD45RO^+^ cultured under iT_EFF_ (open circle), iT_EFF_ + PSGL-1 agonist (open square), or iT_EX_ (open triangle) conditions. Each dot represents a unique donor (5 in total). Data are from 3 independent experiments. Lines are connecting results from the same donor. Bottom: Dot plots showing the retention of IFNγ and TNFα double-producing CD8^+^CD45RO^+^ cultured under iT_EFF_ + PSGL-1 agonist (open square) or iT_EX_ (open triangle) conditions relative to iT_EFF_ (open circle) of the same donor for the 4 out of 5 donors which showed reduction of cytokine production upon PSGL-1 ligation. For each donor, iT_EFF_ relative frequency was set to 100%. Each dot represents a unique donor. Data are from 3 independent experiments. Lines are connecting results from the same donor. (C) Dot plot/bar graph showing the ratio of EOMES/T-bet expression in CD8^+^CD45RO^+^ cultured under iT_EFF_ + PSGL-1 agonist (open square), iT_EX_ (open triangle), or iT_EX_ + PSGL-1 agonist (open diamond) conditions relative to iT_EFF_ (open circle) of the same donor for the 4 out of 5 donors which showed reduction of cytokine production upon PSGL-1 ligation. Ratio was calculated using the relative median florescence intensity (MFI) of Eomes^+^ and/or T-bet^+^ cells. Each dot represents a unique donor. Data are from 3 independent experiments. C57BL/6 mice were infected with LCMV Cl13 and treated I.P. with 100 μg of recombinant human Fc protein (control Fc) or recombinant PSGL-1-human Fc protein (rPSGL-1 Fc) on days 0, 3 and 6 post-infection. NP_(396-404)_ virus-specific CD8^+^ T cell responses were assessed on day 8 post-infection. (D) Top: Representative FACS plots of NP_(396-404)_- tetramer^+^ CD8^+^ T cells from the spleens of LCMV Cl13 infected mice. Bottom: Representative FACS plots of PD-1 and TIM-3 expression within NP_(396-404)_-tetramer^+^ CD8^+^ T cells from the spleens of LCMV Cl13 infected mice. (E) Pie graphs representing the co-expression of inhibitory receptors (PD-1, LAG3, TIM-3) in NP_(396-404)_-tetramer^+^ CD8^+^ T cells from LCMV Cl13 infected mice treated with either control Fc or rPSGL-1 Fc. Data assessed via Boolean gating. C57BL/6 mice were inoculated with 5×10^4^ YUMM1.5 melanoma tumor cells. Beginning on day 0, mice were treated 3x per week I.P. with 100 μg of control Fc or rPSGL-1 Fc. Tumor volume was measured blindly twice per week. (F). Average tumor growth (volume) in YUMM1.5 tumor-bearing mice treated with either control Fc or rPSGL-1 Fc protein. (G) Representative H+E histology combined with anti-CD3 staining in YUMM1.5 tumor sections from control Fc and rPSGL-1 Fc treated mice collected on day 21 post-tumor inoculation. Data in F and G are representative of two independent experiments with 3-5 mice per group in each experiment.

Our studies strongly support the hypothesis that targeting PSGL-1 by immune checkpoint blockade offers a therapeutic approach to promote CD8^+^ T cell responses under conditions of chronic antigen stimulation. Since the available anti-PSGL-1 antibody (clone 4RA10) acts as an agonist, we reasoned that it may have the capacity for cross-linking PSGL-1 on T cells via Fc receptor binding on APCs. Thus, monovalent Fabs were generated and used to treat mice infected with LCMV Cl13. At 30 days post-infection, treatment with Fab’ antibodies did not influence the frequency of NP_(396-404)_-tetramer^+^ virus-specific CD8^+^ T cells in the blood (Extended Data Figure 6A); however, we observed significantly more IFNγ or TNFα SP as well as DP (Extended Data Figure 6B) cytokine-producing CD8^+^ T cells following restimulation with NP_(396-401)_ at day 8 post-infection. A significant increase in IFNγ-producing CD8^+^ T cells was also observed in 4RA10 Fab’ treated mice in response to GP_(33-41)_ peptide stimulation 8 days post-infection (Extended Data Figure 6C). In addition, we noted that co-expression of the inhibitory receptors PD-1, LAG-3, and TIM-3 was decreased in the 4RA10 Fab’-treated group compared to the control group (Extended Data Figure 6D-E).

Since Fab’s by nature have a limited half-life, thus necessitating repeated dosing which could account for the modest limiting effect we observed on T cell exhaustion, we generated a recombinant PSGL-1 with the Fc of human IgG1 (rPSGL-1 Fc). LCMV Cl13 infected mice were treated with rPSGL-1 Fc or recombinant human IgG1 Fc (rIg Fc) as a control and virus-specific CD8^+^ T cell responses were assessed at 8 days post-infection.

The frequency of NP _(396-404)_^+^ CD8^+^ T cells in the spleens was comparable between rIg Fc and rPSGL-1 Fc treated mice but the frequencies of PD-1^+^ and TIM-3^+^ double-positive cells were significantly diminished (Figure 7D). Further, overall co-expression of inhibitory receptors was reduced, consistent with reduced levels of exhaustion (3 inhibitory receptors: 80.3% vs 64.5%)(Figure 7E). Similar results were observed when WT (CD45.2^+^) mice that received WT P14 (CD45.1^+^) CD8^+^ T cells were treated with rIg Fc or rPSGL-1 Fc. We additionally observed significantly decreased TOX expression on a per-cell basis (Extended Data Figure 6F) and increased IFNγ production on a per-cell basis (Extended Data Figure 6G) in mice treated with rPSGL-1 Fc.

Finally, to address whether rPSGL-1 Fc could promote T cell responses to cancer, we evaluated YUMM1.5 tumor growth in mice administered rPSGL-1 Fc or rIg Fc control at the time of tumor inoculation. Strikingly, treatment with rPSGL-1 Fc resulted in significant reduced tumor growth (Figure 7F) that was comparable to that achieved in PSGL-1 deficient mice^18^. Importantly for therapeutic design, we also observed decreased tumor growth when YUMM1.5 tumor-bearing mice received a delayed treatment with rPSGL-1 Fc beginning at day 14 post-tumor injection (Extended Data Figure 6H). T cell infiltration assessed by IHC and anti-CD3 staining indicated that rPSGL-1 treatment enhanced the infiltration and/or expansion of T cells within the tumors (Figure 7G). The Fab’ and rPSGL-1 Fc studies in LCMV Cl13 and the YUMM1.5 melanoma tumor models demonstrate the feasibility of therapeutically blocking PSGL-1 signaling. Together with our new findings identifying the modulation of TCR signaling, T cell metabolism, and the shift in TCF-1 and TOX expression by PSGL-1 signaling, our study here demonstrates the effectiveness in targeting PSGL-1 to relieve immune inhibition of CD8^+^ T cell responses and limit the development of terminal T cell exhaustion.

## Discussion

In this study, we investigated cellular and molecular mechanisms that contribute to the function of PSGL-1 as an intrinsic T cell checkpoint inhibitor. We show that PSGL-1 limits responses to TCR engagement that promote T cell activation and development of T_EFF_ by inhibiting proximal TCR signaling via Zap70 as well as downstream activation of signaling via Erk1/2 and Akt. By phosphoproteomics, we identified that with PSGL-1 deficiency, multiple proteins involved in T cell signaling and activation display increased phosphorylation in response to TCR signaling, demonstrating a broad role for PSGL-1 in the regulation of T cell activation. It is notable that PSGL-1 inhibits T cell activation with limited TCR engagement and with low affinity TCR ligands, indicating that PSGL-1 is a fundamental regulator that plays a role in preventing excessive T cell responses. The regulatory function of PSGL-1 could be attributed to decreased expression of an inhibitor of T cell activation, Sts-1, an E3 ubiquitin ligase that targets Zap70 for degradation^20,43^. This protein is encoded by the gene, *Ubash3b*, whose expression we found to be greatly diminished in both T_EX_ and T_EFF_ with PSGL-1 deficiency by RNA-sequencing. We also detected closed chromatin for an enhancer region of *Ubash3b*, and confirmed decreased Sts1 protein expression in PSGL-1 deficient T cells. Deficiency of Sts-1 and 2 proteins was previously shown to lead to hyperreactive TCR signaling in mice, and increased susceptibility to autoimmunity, although only Sts-1 was shown to regulate Zap70^20^. Our finding that Sts-1 is diminished but not absent in PSGL-1^-/-^ T cells suggests that TCR signaling is enhanced without driving terminal T cell differentiation.

Mechanistically, Zap70 in T cells is a homologue to Syk which has been shown in neutrophils to engage PSGL-1 signaling through ERM proteins that regulate its migration to the uropod of the cells via actin rearrangements^44^. By microscopy, we show that PSGL-1 and CD3 of the TCR complex can colocalize on the T cell surface when only CD3 is engaged, but that colocalization is much more compact when both molecules are engaged. These findings indicate that although PSGL-1 can be redistributed to the uropod of migrating cells, on T cells, PSGL-1 is localized with the TCR where it can contribute to the strength of early T cell signaling^45^. We hypothesize that in this context, Sts-1 limits TCR signaling through its ubiquitination function that targets Zap70 for degradation. As a result, we hypothesize that in the absence of PSGL-1, reduced Sts-1 expression supports enhanced T cell signaling to promote the development of effector functions which rely upon the dramatically enhanced glycolytic metabolism and glucose uptake, and to a lesser extent, oxygen consumption, as shown by *in vitro* Seahorse metabolic assays. In addition, from single cell analyses of tumor infiltrating T cells, we found that metabolic reprogramming in PSGL-1 deficient T cells promotes the expression of multiple enzymes that regulate glycolysis at the level of gene transcription, where increased expression was concomitantly associated with the expression of genes involved in effector T cell functions. These findings demonstrate that PSGL-1-dependent regulation of TCR signal strength constrains CD8 T cell metabolic activity to limit anti-tumor responses, providing a mechanism by which PSGL-1 acts as an immune checkpoint inhibitor.

Previous studies support a conclusion that an ability to maintain T glycolytic metabolism in the tumor microenvironment where T cells compete with tumor cells for glucose is a fundamental mechanism by which CD8^+^ T cells can promote more effective antitumor responses^4,46^. However, more broadly, we show that enhanced effector responses of PSGL-1 deficient T cells are accompanied by altered differentiation in the settings of chronic antigen-stimulation in the LCMV Cl13 model where subsets of exhausted CD8^+^ T cells have been defined^28-30^ and where gene signatures of T_EX_ subsets in LCMV Cl13 and B16-OVA melanoma have significant overlap^47^. Most notable was the significantly decreased expression of TOX with the maintenance of TCF-1, a transcriptional profile that predicts decreased CD8^+^ T cell exhaustion. Our data suggest that PSGL-1 is a regulator of terminal T_EX_ differentiation, and that in its absence, CD8 T cells can retain stemness and a capacity to maintain responsiveness. In the B16-OVA model, that is variably responsive to anti-PD-1 treatment, most PSGL-1^-/-^ T cells were double positive for TCF-1 and TOX. It is likely that the extent of T cell responses varies in these different models due to differences in the levels of inflammation, but in all of these settings, much better responses occurred with PSGL-1 deficiency, and terminal T_EX_ differentiation was curtailed. Since we previously found that PSGL-1 deficiency was associated with decreased expression of multiple inhibitory receptors, one possible explanation for the lack of terminal T_EX_ differentiation is reduced engagement of several pathways of immune inhibition^48,49^. However, our studies demonstrate that although ligation of PSGL-1 promotes T cell exhaustion and high expression of inhibitory receptors, PSGL-1 does not directly regulate PD-1 expression. Since a previous study showed that TCR signaling in exhausted T cells is strongly inhibited during chronic LCMV infection, our results support a conclusion that promoting increased responses to TCR engagement to even low affinity ligands is the fundamental mechanism by which exhaustion is curtailed^6^. We propose greater and more sustained TCR signaling and metabolism with PSGL-1 deficiency is of central importance to preventing the differentiation of terminal T_EX_ in melanoma tumors and possibly other cancers.

Our results strongly support the conclusion that PSGL-1 is an important target for immune checkpoint blockade. Our data using a monovalent Fab’ anti-PSGL-1 reagent to block PSGL-1 in the LCMV Cl3 response, and a recombinant PSGL-1 protein to block PSGL-1 in the LCMV Cl13 and YUMM1.5 models validate that ICB can recapitulate many aspects of PSGL-1 deficiency. Further studies will be required to determine whether or how PSGL-1 blockade affects the responses of other PSGL-1 expressing cells in our models as well as to address whether exhaustion in T cells can be reversed after it has become established. Importantly, however, we show that agonism of PSGL-1 signaling in the context of TCR signaling promotes exhaustion in both human and mouse T cell responses, underscoring the conclusion that one of the highly conserved functions of PSGL-1 is that of a negative regulator of T cell activation and responses. Since the LCMV Cl13 and B16 models are not highly sensitive to anti-PD-1 treatment, we propose that PSGL-1 blockade could synergize with anti-PD-1. Although PD-1 blockade can enhance TCR signals ^6^ as well as reduce metabolic insufficiencies of T_EX_^27^, and deficiency of PD-1 can enhance CD8^+^ T cell effector responses to chronic LCMV, its genetic absence can promote the accumulation of terminally exhausted T cells^50^. This outcome, along with the finding that terminal T cell exhaustion is irreversible^51^, suggests that repeated blockade of PD-1 could contribute to acquired resistance of cancer patients to ICB therapy. It is noteworthy in this regard that PSGL-1 deficiency in T cells intrinsically promotes stemness in the settings of both acute and chronic antigen stimulation. Thus, targeting PSGL-1 could enhance the potential of achieving more sustained antitumor T cell responses. In particular for those tumors shown to be highly resistant to PD-1 targeted therapies such as pancreatic and prostate cancers, PSGL-1 blockade could represent a new target for immunotherapy.

## Acknowledgements

The authors would like to thank the invaluable contributions from Dr. Kristen Jepsen and the Institute for Genomic Medicine (IGM) at UC San Diego for their assistance with the sc-RNA sequencing experiments, and Dr. Kathleen Fisch and the UC San Diego Center for Computational Biology & Bioinformatics (CCBB) for their assistance with scRNA-seq data analysis. We thank Petrus de Jong for his assistance with the initial design of the Seahorse metabolism assays. We thank Bobby Ng for his assistance with Western blots. We would like to thank all members of the Bradley lab for their assistance in completing this study. This work was supported in part by the Sanford Burnham Prebys NCI-Designated Cancer Center Support Grant, P30 CA030199. We would like to acknowledge the contributions of the following Sanford Burnham Prebys Core facilities: the Flow Cytometry Core (Yoav Altman and Amy Cortez), the Vivarium Core (Buddy Charbono and Andy Vasquez), the Histology Core (Gia Garcia), the Bioinformatics Core (Jun Yin, Ph.D.), the Proteomics Core (Alex Campos, Ph.D.), and the Cancer Metabolism Core (David Scott, Ph.D.).

## Author Contributions

J. Hope completed the metabolism assays, single-cell RNA-sequencing experiments, tumor studies, LCMV studies, *in vivo* recombinant PSGL-1 studies, and *in vitro* assays. D. Otero completed the T cell signaling studies, microscopy, and *in vitro* assays. J. Hope designed and H. Faso completed human PBMC assays. C. Stairiker, E. Bae, A. Palete, H. Faso, and M. Henriquez completed tissue processing, tumor measurements, *in vitro* assays, flow cytometry staining, and mouse breeding and genotyping. H. Seo and A. Rao completed ATAC-seq experimental design assistance and library preparation. A. Campos performed proteomics/phosphoproteomics and analysis. J. Yin completed ATAC-seq analyses. X. Lie and P. Adams completed ATAC-seq enhancer region comparisons and ATAC-seq interpretation assistance. D. Otero and E. Wang completed Western blots. R. Tinoco completed *in vivo* Fab’ 4RA10 studies. J. Hope, D. Otero, and L. Bradley designed the experiments and completed data analysis. J. Hope and L. Bradley wrote the manuscript. All authors reviewed, critiqued, and approved the manuscript prior to submission.

## Funding

JH is currently supported by an American Cancer Society Postdoctoral Fellowship (PF-20-113-01-LIB) and previously by the Sanford Burnham Prebys Frontiers in Immunology T32 Postdoctoral Fellowship (T32 AI125209). EB was supported through The American Association of Immunologists Careers in Immunology Fellowship Program. These studies were supported in part by the following grants: awarded to LB -R01 AI106895, R21 CA249353, R21 CA216678, R03 CA252144, Melanoma Research Alliance MRA 696326, Department of Defense W81XWH-20-1-0324; awarded to AR – AI140127 and AI109842. This work was supported in part by the Sanford Burnham Prebys NCI-Designated Cancer Center Support Grant, P30 CA030199.

## Declaration of Interests

L. Bradley and R. Tinoco hold the patent for targeting PSGL-1 on T cells in the treatment of infectious diseases, cancers, and immune and inflammatory diseases.

## Methods

### Mice

C57BL/6J, B6.Cg-PSGL-1^*tm1Fur*^/J (PSGL-1^*-/-*^), and C57BL/6 Tg(TcraTcrb)1100Mjb/J (OT-I) were obtained from the Jackson Laboratory. PSGL-1^*-/-*^ were backcrossed to the Thy1.1 background (B6.PL-*Thy1*^*a*^*/*CyJ), and further crossed with OT-I mice to establish these lines with PSGL-1 deficiency. All mice were kept in a barrier facility (certified by the Association for the Assessment and Accreditation of Laboratory Animal Care) on acidic water at Sanford Burnham Prebys Medical Discovery Research Institute. After LCMV infection, mice are maintained in BSL2 containment. All lines in the Bradley mouse colony are backcrossed to parental C57BL/6J or B6.PL-*Thy1*^*a*^*/*CyJ lines annually. This study was carried out in accordance with the recommendations and approval of the Institutional Animal Care and Use Committee (IACUC).

### Tissue Culture

The YUMM1.5 melanoma tumor cell line were cultured in Iscove’s Modified DMEM (IMEM) supplemented with 10% fetal bovine serum (FBS), 10^5^ I.U. Penicillin, 10^5^ μg/mL Streptomycin, and 292 mg L-glutamine (Corning Cellgro). B16-OVA melanoma tumor cells were cultured in DMEM supplemented with 10% fetal bovine serum (FBS, Corning), 10^5^ I.U. Penicillin, 10^5^ μg/mL Streptomycin, 292 mg L-glutamine, and 2 μg/mL Geneticin. Cells were maintained *in vitro* at 37°C, 5% CO_2_. For *in vivo* experiments, tumor cell aliquots from the same passage were thawed one week and passaged 3 times prior to injection. For culturing of human and mouse T cells, cells were cultured in complete T cell media: RPMI 1640 supplemented with 10% FBS, 10^5^ I.U. Penicillin (Gibco), 10^5^ μg/mL Streptomycin (Gibco), [10 mM] HEPES buffer (Gibco), [1 mM] Sodium Pyruvate (Gibco), 1X MEM nonessential amino acids (Gibco), and [0.55 mM] betamercaptoethanol (Gibco).

### Infection and Tumor Studies

For tumor initiation, 6–8-week-old male or female mice were anesthetized with 2.5% isofluorane gas, their hind flank was shaved, and tumor cells (resuspended in Matrigel (Corning):1X PBS at a 1:1 ratio) were injected subcutaneously (s.c.) in a volume of 100 μL PBS. For adoptive cell transfers, recipient mice received a tail vein injection of T cells in a volume of 100 μL PBS as indicated for the individual experiments. When naïve T cells were transferred, cells were first mixed 1:10 with C57BL/6 splenocytes obtained from uninfected/non-tumor-bearing wildtype mice. Tumor growth was assessed three times per week. For YUMM1.5 studies, only male mice were used due to the presence of male antigen on the YUMM1.5 tumors. At the indicated time points, tumors, spleens, and inguinal draining and non-draining lymph nodes were collected. For LCMV Clone 13 infection, 6–8-week-old male and female mice received an intravenous injection of 2.5×10^6^ FFU in a final volume of 100 μL PBS. At the indicated time points, spleens were collected.

### Tissue Processing

Tumors were processed using the Miltenyi Mouse Tumor Dissociation Kit and gentleMACS C tubes (Miltenyi) according to manufacturer’s instructions, except with the intentional exclusion of enzyme R (due to cleavage of PSGL-1 and CD44). Spleens and lymph nodes were processed into single-cell suspensions by manual dissociation through a 70 μM filter (Falcon) using a cell strainer pestle (CELLTREAT). Tumors, spleens, and lymph nodes were processed and maintained in RPMI 1640 (Corning) supplemented with 5% FBS, 1% Penicilin/Streptamycin/L-glutathione. Tumors and spleens were treated with RBC lysis buffer (0.145 M NH_4_Cl, 0.05 M Tris-HCl, pH 7.2) for ∼1 minute at room temperature.

### Naïve T cell enrichment

For all experiments in which isolated CD8^+^ T cells were used, untouched naïve (CD44^-^CD62L^+^) CD8^+^ T cells were enriched via negative selection using biotinylated antibodies at the concentration indicated in Table 1 and using MojoSort Streptavidin Nanobeads (BioLegend). Enrichments were completed in 1X HBSS (Ca^2+^ and Mg^2+^ free) containing 2% FBS and 1 mM EDTA at a cell concentration of 1×10^8^/mL. CD8^+^ T cells enriched in this manner were on average >98% pure and depleted of CD44-expressing CD8^+^ T cells. In all experiments, naïve CD8^+^ T cells were enriched from 6-8 week old male or female mice (depending on the experiment and matching the host recipients in the case of adoptive T cell transfers).

### T Cell Signaling Assays

Naïve CD8^+^ T cells were enriched from WT or PSGL-1^-/-^ splenocytes as above. After enrichment, 1×10^6^ naïve OT-I CD8^+^ T cells were activated with immobilized anti-CD3ε (145-2C11, BioXcell) in 24-well plates at the anti-CD3ε concentration or for the time indicated. For analysis of activation status, cells were stained for assessment by flow cytometry as described below. For Western blot analysis of phosphorylated and total expression levels of signaling molecules, activated cells (and non-activated controls) were washed twice with 1X HBSS and 3-5×10^6^ cells were pelleted for lysis in mammalian protein extraction reagent (M-PER, Fisher). Protein concentration was normalized for loading based on a bicinchoninic acid (BCA) assay and mixed with NuPage LDS Sample Buffer (Life Technologies) before loading. Samples were run on a 4-12% Bis-Tris gradient gel (Life Technologies) using the XCell Mini-cell electrophoresis system in 1X MES SDS Running Buffer (Life Technologies). Proteins were transferred to a nitrocellulose membrane using the XCell II Blot Module and 1X Bolt Transfer Buffer with 20% methanol (Life Technologies). Blots were blocked with blocking buffer (LICOR) then incubated overnight with the indicated primary antibody, washed with TBS-T (Cell Signaling), and incubated for 1 hour with appropriate secondary antibodies in blocking buffer (LICOR). Blots were imaged using a LICOR Odyssey and quantified using ImageJ. For phosflow flow cytometry analysis of phosphorylated signaling molecules, naïve WT (CD45.1^+^) and PSGL-1^-/-^ (CD90.1^+^) OT-I CD8^+^ T cells were mixed 1:1 with each other and then with C57BL/6 splenocytes generated from uninfected female mice. T cells were activated in FACS tubes (BD Falcon) by the addition of 10 μM SIINFEKL peptide (GenScript) for the indicated time at 37°C. After activation, pre-warmed 1X BD Phosflow Fix Buffer I (BD) was immediately added to cells and incubated for 15 minutes at 37°C. Cells were washed with BD Phosflow Perm Wash and permeabilized by the addition of cold BD Phosflow Perm Buffer III while vortexing. After permeabilization, both surface markers and phosphorylated signaling molecules were stained simultaneously in BD Phosflow Perm Wash for 45 minutes. Cells were washed and fixed with 1% formaldehyde prior to data acquisition on a BD LSRFortessa X-20 (BD) and analyzed using FlowJo v10 software (BD).

### STS-1 Western blot analysis

For STS-1 Western blot analysis, cells were lysed in M-PER buffer (Thermo Scientific) containing protease/phosphatase inhibitor cocktail (Roche). Protein concentration was measured using a BCA assay (Pierce). Equivalent amounts of each sample were loaded on 4-12% Bis-Tris gels (Invitrogen), transferred to nitrocellulose membranes, and immunoblotted with antibodies against STS-1 (Proteintech #19563-1-AP) and β-Actin (Cell Signaling #3700). IRDye®800-labeled goat anti-rabbit IgG and IRDye®680-labeled goat anti-mouse IgG (LI-COR) secondary antibodies were used and detected on an Odyssey CL_x_ system.

### Mouse *In vitro* T Cell Exhaustion

Cells were cultured in complete T cell media supplemented with 5 ng/mL each of recombinant murine IL-7 and IL-15 (PeproTech). One day prior to activation of OT-I cells, C57BL/6 splenocytes were made into a single-cell suspension and treated overnight with IFNγ(5 ng/mL) to induce expression of CD80 and CD86 on antigen presenting cells. Splenic OT-I and PSGL-1^*-/-*^ OT-I CD8^+^ T cells were isolated by negative selection with magnetic beads from uninfected female mice 8-10 weeks of age. Cells were cultured at a 1:1 ratio of CD8^+^ T cells and IFNγ-treated splenocytes at 1 × 10^6^/mL with either one-time stimulation with 50 ng/mL OVA_(257-264)_ peptide for 48 hours (single stim), or with daily peptide stimulation with 50 ng/mL of OVA_(257-264)_ peptide for 5 days (repeated stim). In experiments using SIINFEKL variant peptides, cells received either a one-time stimulation or daily peptide stimulation with 50 ng/mL of the indicated peptide. For all conditions, after 48 hours of culture, cells were washed twice and re-seeded at 0.5×10^6^ cells/mL in complete T cell media. On days two and five, all conditions were harvested and the cells were counted. CD8^+^ T cells were stained as described below (flow cytometry) to measure activation marker, inhibitory receptor, and transcription factor expression and cytokine production following restimulation with OVA_(257-264)_.

### Human *In vitro* T Cell Exhaustion

Cells were cultured in complete T cell media supplemented with 5 ng/mL each of recombinant human IL-7 and IL-15 (PeproTech) and 10 U/mL recombinant human IL-2. Cryopreserved healthy donor human PBMCs (purchased from iXcells Biotech) were thawed then rested for five hours at 37°C. After resting, cells were cultured at a concentration of 1 × 10^6^ CD3^+^ cells/mL and stimulated with 6.25 μL/mL of ImmunoCult Human CD3/CD28 T cell activator (StemCell). For PSGL-1 agonist, tissue culture plates were coated with 5 μg/mL of anti-human CD162 (KPL, Biolegend). On day three of activation, cells were washed and re-seeded at 1×10^6^ cells/mL. Cells were then divided into either iT_EFF_ or iT_EX_ conditions. iT_EFF_ conditions did not receive additional ImmunoCult after the initial 72 hours; iT_EX_ conditions received an additional 6.25 μL/mL of ImmunoCult each day on days three through eight. All conditions were harvested, washed and re-seeded as above on day 6. On day 9, T cells were stained for assessment by flow cytometry as described below to measure activation marker, inhibitory receptor, and transcription factor expression and cytokine production following restimulation with ImmunoCult overnight in the presence of Brefeldin A and monensin.

### 2-NBDG Uptake Assays

All solutions were prepared in glucose-free, serum-free RPMI. 2-NBDG (Cayman Chemical) uptake was performed with a 200 μM solution. Cells were incubated at 37°C for 30 minutes before proceeding with FACS staining. Cells were resuspended in FACS wash and analyzed within 30 minutes of completion of FACS staining.

### Flow Cytometry

In all stains, cells were pre-treated with 2.5 μg/mL anti-CD16/32 (Fc Block; 2.4G2; BioLegend, San Diego, CA) for 15 minutes before continuing with surface staining. For surface stains, 2×10^6^ cells were stained for 20 min on ice. Cells were stained with the fluorochrome conjugated monoclonal antibodies as indicated (Extended Data Table 1). Where indicated, cells were also stained with APC labeled-tetramers of H-2^b^ major histocompatibility complex class I loaded with GP_(33-41)_, or BV421 labeled-tetramers of H-2^b^ major histocompatibility complex class I loaded with NP_(396-404)_ provided by the NIH Tetramer Core Facility. After staining, cells were washed twice with 1X HBSS containing 3% FBS and 0.02% sodium azide and fixed with 1% formaldehyde. For staining of intracellular transcription factors, 2×10^6^ cells were stained as above for surface markers. Following the surface stain, cells were permeabilized with eBioscience FoxP3 fixation buffer (ThermoFisher) overnight at 4°C, washed using eBioscience 1X Perm Wash (ThermoFisher), followed by intracellular staining for 45 minutes. Cells were then washed 2 times with 1X Perm Wash containing 0.02% sodium azide and fixed with 1% formaldehyde. Cells were stained with the fluorochrome conjugated monoclonal antibodies indicated in the table below. For staining of intracellular cytokines, 2×10^6^ cells were stimulated with the indicated peptide (OVA: 5 μg/mL SIINFEKL peptide; GP_(33-41)_: 2 μg/mL KAVYNFATM peptide; NP_(396-404)_: 2 μg/mL FQPQNGQFI; all from Genscript) or ImmunoCult (for human PBMCs, StemCell) overnight at 37°C, 5% CO_2_ in the presence of GolgiPlug (BD Biosciences), Monensin (BioLegend), and 10 U/mL recombinant human IL-2. Cells were surface stained as above then fixed overnight at 4°C with FoxP3 fixation buffer (eBioscience), washed using Perm/Wash buffer (eBioscience) and stained for intracellular cytokines for 45 minutes at 4°C. Cells were stained with the fluorochrome conjugated monoclonal antibodies indicated in the table below. 1X Perm Wash containing 0.02% sodium azide and fixed with 1% formaldehyde. All data was acquired on a BD LSRFortessa X-20 (BD) and analyzed using FlowJo v10 software (BD).

### Fluorescence Microscopy and Amnis Imaging Flow Cytometry

Naïve OT-I CD8^+^ T cells were enriched from WT or PSGL-1^-/-^ splenocytes from 6-8 week old female mice using magnetic bead enrichment. After enrichment, a subset of cells was directly fixed with 1% formaldehyde. For crosslinking, naïve cells were incubated with anti-CD3ε (145-2C11, BioXcell), anti-PSGL-1 (4RA10, BioXcell), or both for 15 minutes at 4°C. After washing, cells were crosslinked with biotinylated anti-hamster IgG (for anti-CD3; BD Pharmingen) and/or Goat anti-rat IgG (for anti-PSGL-1, Sigma Aldrich) and incubated for 10 minutes at 37°C. After incubation, cells were fixed as above. CD3 localization was detected using FITC conjugated streptavidin (CALTAG Laboratories) and PSGL-1 localization was detected using anti-Rat IgG-PE (BioLegend) following a 30 minute incubation at 4°C. Cells were plated on slides pre-coated with CellTak (Corning) for analysis by fluorescence microscopy. For quantification by Amnis Imaging Flow Cytometry, samples were acquired using an ImageStreamX Mk11 (2 cameras) and INSPIRE v200.1.388.0 software. Images were assessed at a 40X objective in high resolution/low speed mode. 10,000 gated events (Hoechst+FITC+, in focus, single cells) were acquired per condition. For analysis, IDEAS v6.2.183.0 software was used and compensation and nuclear localization were calculated using the IDEAS wizard. The Amnis similarity score (a transformed Pearson’s correlation coefficient ^52^) was calculated from these data.

### Seahorse

Assays were conducted in Seahorse XF RPMI Medium, pH 7.4 (Agilent) supplemented with sodium pyruvate [1 mM], L-glutamine [2 mM], and glucose [5 mM]. Seahorse XF plates were pre-coated with 22.4 ug/mL of Cell-Tak solution (Corning) as per manufacturer’s instructions on the same day as the assay and 0.3×10^6^ cells were seeded per well. For glycolytic rate assays: rotenone [0.0013 mM]/antimycin A [0.013 mM], and 2-DG [500 mM] prepared in supplemented Seahorse XF RPMI Medium were injected sequentially as indicated. Assays were conducted using an XFp (8 well) system and analyzed using WAVE software (Agilent).

### Single-Cell RNA-sequencing

Naïve WT (CD45.1^+^) or PSGL-1^*-/-*^ (CD90.1^+^) OT-I CD8^+^ T cells were activated with immobilized anti-CD3ε (145-2C11, BioXcell) and anti-CD28 (37.51, BioXcell)(5 μg/mL, each) for 3 days prior to transfer. 1×10^6^ activated WT or PSGL-1^-/-^ OT-I CD8^+^ T cells were then injected I.V. into tumor-bearing C57BL/6 (CD45.2^+^/CD90.2^+^) mice, 7 days after inoculation with B16-OVA tumors. Three days after T cell transfer, live donor OT-I CD8^+^ T cells were sorted by FACS (fluorescence activated cell sorting) from the tumor-draining and non-draining inguinal lymph nodes in a subset of mice. Six days after T cell transfer, live donor OT-I CD8^+^ T cells were sorted by FACS from the tumor-draining and B16-OVA tumors in a subset of mice. For both data sets, live sorted cells were immediately processed on a 10X Chromium (10X) to generate single cell 3’ libraries using the 10X Single Cell 3’ Reagent Kits v2 as per manufacturer’s instructions (Rev E, 2018). Data were acquired using a HiSeq 400 PE50 at the Institute for Genomic Medicine Facility at the University of California, San Diego.

### scRNA-seq Analysis

scRNA-seq data were aligned, counted, and aggregated using Cell Ranger v2.2.0 software (10X Genomics). CellLoupe (10X Genomics) was used to visualize data. SeqGeq (FlowJo, BD) was used in some analyses to assess expression changes within subgated “cell” populations. Data quality control was assessed using Seurat based on Satija Lab tutorials and was completed by the Center for Computational Biology and Bioinformatics center at the University of California, San Diego. Heatmaps of the top 10 most differentially expressed genes within each cluster were generated using Seurat “DoHeatmap”.

### Proteomics and Phosphoproteomics Sample Preparation

5×10^6^ naïve CD8^+^ T cells were plated per well of a 6-well plate coated with 5 μg/mL anti-CD3ε antibody (145-2C11, BioXCell)(activated) or appropriate IgG control (naïve) and centrifuged for 1 minute at 300xg. Per biological replicate, cells were pooled from 2-3 individual mice. Cells were incubated for 15 minutes at 37°C, then placed directly on ice for removal by pipetting. Cell pellets were snap frozen at -80°C then lysed in 8M urea, 50 mM ammonium bicarbonate (ABC) and Benzonase. Cellular debris were removed and supernatant protein concentration was determined using a BCA protein assay (Thermo Scientific). Disulfide bridges were reduced with 5 mM tris(2-carboxyethyl)phosphine (TCEP) at 30°C for 60 min, and cysteines were subsequently alkylated with 15 mM iodoacetamide (IAA) in the dark at room temperature for 30 min. Urea was then diluted to 1 M urea using 50 mM ABC, and proteins were subjected to overnight digestion with mass spec grade Trypsin/Lys-C mix (Promega). Following digestion, samples were acidified with formic acid (FA) and subsequently peptides were desalted using AssayMap C18 cartridges mounted on an AssayMap Bravo Platform (Agilent Technologies). Two biological replicates were pooled together to generate 100 micrograms of total peptide for TMT labeling and subsequent fractionation and phosphopeptide enrichment. Dried pooled sample was reconstituted in 20 mM ammonium formate pH ∼10, and separated in 2 aliquots of 100 and 900 micrograms for total proteomics and phosphoproteomics fractionation. Total proteome TMT and phosphoproteome aliquots were fractionated using a Waters Acquity BEH C18 column (Total: 2.1x 15 cm, 1.7 μm pore size; Phospho: 4.6x 25 cm, 3.5 μm pore size) mounted on an M-Class Ultra Performance Liquid Chromatography (UPLC) system (Waters)(total) or Vanquish Horizon UPLC system (ThermoFisher)(phospho). A total of 36 (total) or 12 (phospho) fractions were collected and pooled in a non-contiguous manner into 18 total fractions and dried to completeness in a SpeedVac concentrator prior to mass spectrometry analysis. TMT-labeled phosphopeptides were enriched in an automated fashion using the AssayMAP Bravo Platform (Agilent Technologies).

### LC-MS/MS analysis

Dried peptide fractions were reconstituted with 2% ACN, 0.1% FA and analyzed by LC-MS/MS using a Proxeon EASY nanoLC system (Thermo Fisher Scientific) coupled to an Orbitrap Fusion Lumos mass spectrometer (Thermo Fisher Scientific). Peptides were separated using an analytical C18 Aurora column (75μm x 250 mm, 1.6μm particles; IonOpticks) at a flow rate of 300 nL/min using a 75-min gradient: 1% to 6% B in 1 min, 6% to 23% B in 44 min, 23% to 34% B in 28 min, and 27% to 48% B in 2 min (A= FA 0.1%; B=80% ACN: 0.1% FA). The mass spectrometer was operated in positive data-dependent acquisition mode. MS1 spectra were measured in the Orbitrap with a resolution of 60,000, at accumulation gain control (AGC) target of 4e5 with maximum injection time of 50 ms, and within a mass range from 350 to 1500 m/z. Tandem MS was performed on the most abundant precursors with charge state between +2 and +7 by isolating them in the quadrupole with an isolation window of 0.7 m/z. Precursors were fragmented with higher-energy collisional dissociation (HCD) with normalized collision energy of 35% and the resulting fragments were detected in the Orbitrap at 50,000 resolution, at AGC of 1e5 and maximum injection time of 105 ms. The dynamic exclusion was set to 20 sec with a 10 ppm mass tolerance around the precursor. To increase confidence of the data, each fraction of total and enriched phosphopeptide were run in duplicates. All mass spectra were analyzed with MaxQuant software version 1.6.11.0. MS/MS spectra were searched against the Mus musculus Uniprot protein sequence database (downloaded in January 2020) and GPM cRAP sequences (commonly known protein contaminants). Reporter ion MS2 type was selected along with TMT 10plex option. Precursor mass tolerance was set to 20ppm and 4.5ppm for the first search where initial mass recalibration was completed and for the main search, respectively. Product ions were searched with a mass tolerance 0.5 Da. The maximum precursor ion charge state used for searching was 7. Carbamidomethylation of cysteine was searched as a fixed modification, while oxidation of methionine and acetylation of protein N-terminal were searched as variable modifications. Enzyme was set to trypsin in a specific mode and a maximum of two missed cleavages was allowed for searching. The target-decoy-based false discovery rate (FDR) filter for spectrum and protein identification was set to 1%. Statistical analysis of TMT total and phosphoproteome data were carried out using in-house R script (version 3.5.1, 64-bit), including R Bioconductor packages. First, TMT reporter intensities were log2-transformed and normalized (loess normalization) across samples to account for systematic errors. Following normalization, all non-razor peptide sequences and precursor isolation interference (MaxQuant “PIF” column in evidence file) below 0.8 (total proteome) or 0.9 (phosphoproteome) were removed from the list. Phosphopeptide-and Protein-level quantification and statistical testing for differential abundance were performed using MSstatsTMT bioconductor package ^53^. In MSstatsTMT experimental setting, the technical replicates were analyzed together as different ‘mixtures’.

### ATAC-sequencing

Naïve OT-I CD8^+^ T cells were enriched from WT or PSGL-1^-/-^ splenocytes from 6-8 week old female mice using magnetic bead enrichment. After enrichment, 1×10^6^ naïve CD8^+^ T cells were plated per well of a 24-well plate coated with 5 ug/mL anti-CD3 antibody (145-2C11, BioXCell) and centrifuged for 1 minute at 300xg. Cells were incubated for 2 hours at 37°C. ATAC-seq libraries were prepared following the omni-ATAC protocol^54^ with minor changes. 50,000 cells were rinsed with 1XPBS and centrifuged at 400g for 3 minutes. Cells were resuspended in 50 μl lysis buffer (10 mM 1 M Tris-HCl pH 7.4, 10 mM NaCl, 3 mM MgCl^2^, 0.1% NP40, 0.1% Tween 20, 0.01% Digitonin), and incubated at 4°C for 5 minutes, after which 1 mL rinsing buffer (10 mM Tris-HCl pH 7.4, 10 mM NaCl, 3 mM MgCl^2^, 0.1% Tween 20) was added, then centrifuged at 1,000xg for 10 minutes at 4°C. After removing the supernatant, the nuclei were resuspended in 50 μL of transposition mix (25 μL of TD buffer [20 mM Tris-HCl pH 7.6, 10 mM MgCl2, 20% Dimethyl Formamide], 2.5 μL of 2 μM transposase 16.5 ml PBS, 0.5 μL 1% digitonin, 0.5 μL 10% Tween-20, 5 mL water) and incubated at 37 °C for 30 minutes. DNA was obtained with a Qiagen MinElute Kit (Qiagen). Libraries were prepared with KAPA HiFi High sensitivity Real-time PCR master mix. The ATAC library sequenced on an Illumina Novaseq 6000 sequencer (paired-end 50-bp reads) at the La Jolla Institute for Immunology sequencing core.

### ATAC-seq Analysis

ATAC-seq QC and analysis was performed by the Bioinformatics Core at Sanford Burnham Prebys Medical Discovery Institute. In brief, FASTQ files were generated following adaptor trimming (Cutadapt) and MultiQC was used to evaluate library quality. STAR was used to align ATAC-seq data to mouse genome B38, Ensembl v84 and Homer was used for peak calling and annotation. DESeq2 and Homer were used to identify differential peak analysis. IGV genome browser was used for peak visualization. Reproducible peaks were compared with public dataset [PMID: 28288100] using BEDTools ^55^.

### Statistics

Specific statistical analyses for big data sets are indicated in their specific methods section. For flow cytometry data, the normality of the population distribution was assessed using the Shapiro-Wilk normality test by GraphPad Prism 9. Significant differences between normally distributed populations were assessed using a two-tailed, unpaired *t*-test; significant differences between non-normally distributed populations were assessed using a two-tailed Mann Whitney exact test. The tests performed are denoted in each figure legend and subsequent p-values are annotated in the associated figure.

## Notes

### Competing Interest Statement

The authors have declared no competing interest.

