## Extended Data for "PSGL-1 directs early TCR signaling to repress metabolism and promote T cell exhaustion by modulating the TCF-1/TOX axis in CD8^+^ T cells"

**Extended Data Figure 1.**

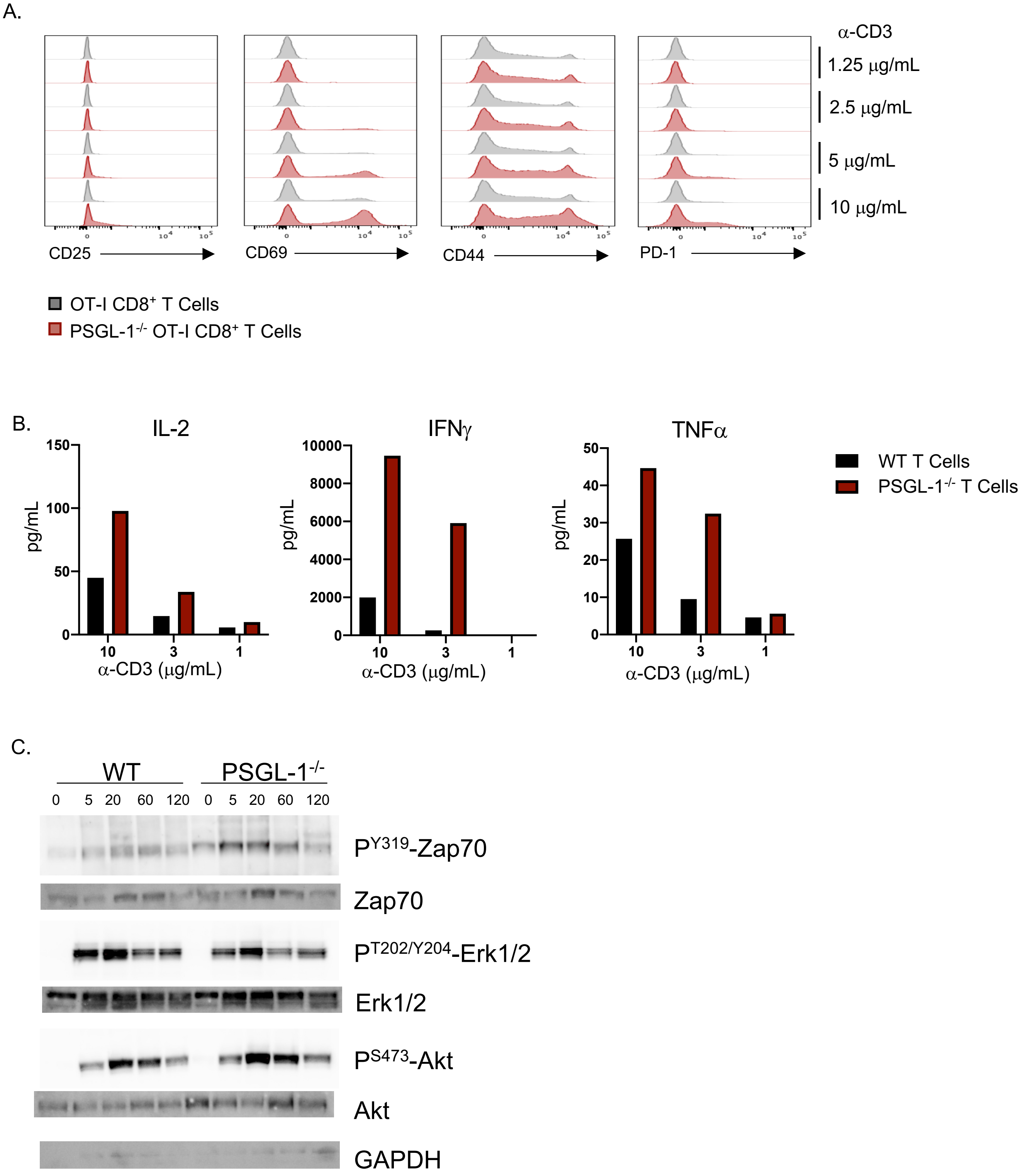

**Extended Data Figure 2.**

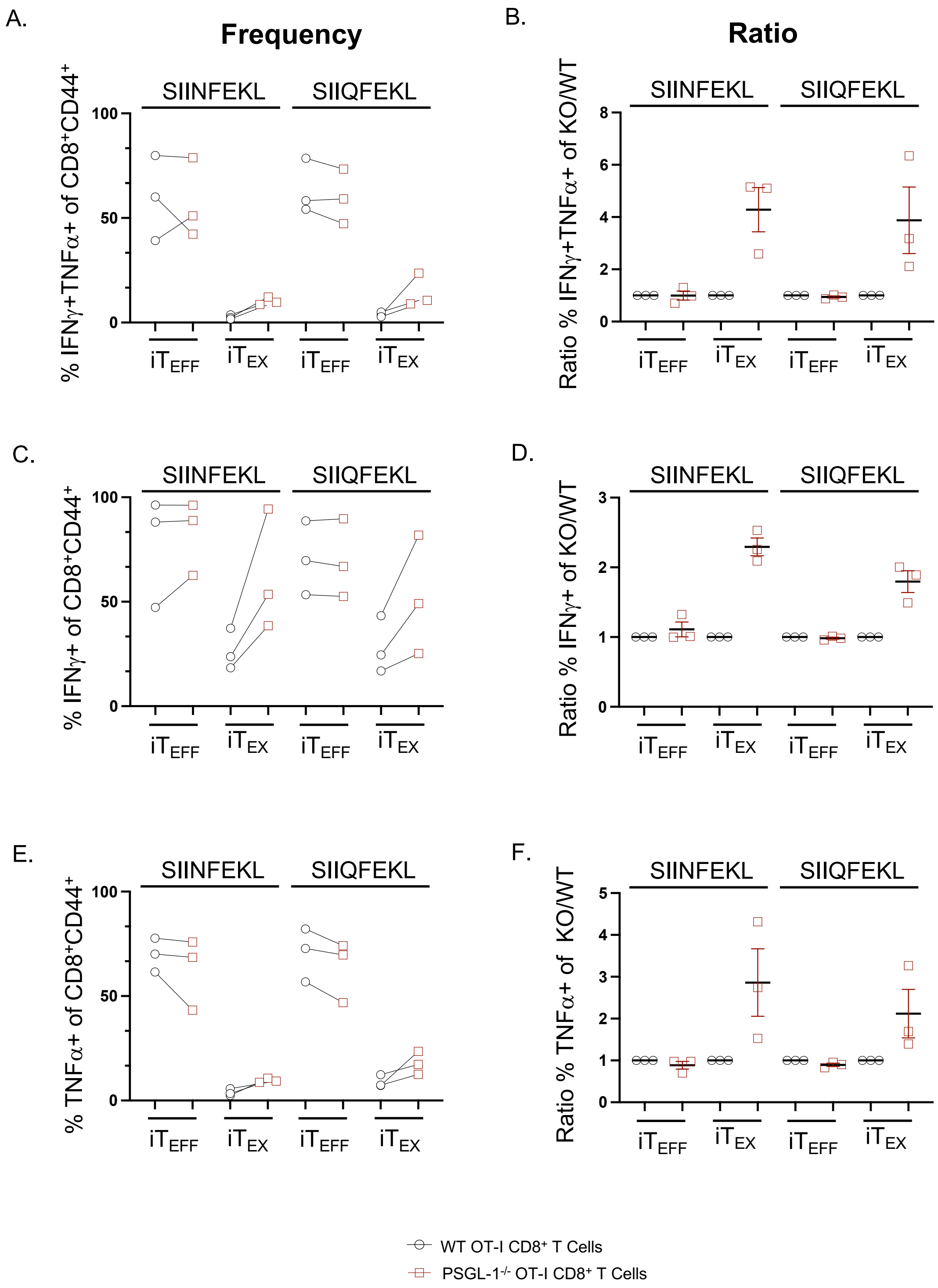

**Extended Data Figure 3.**

A.

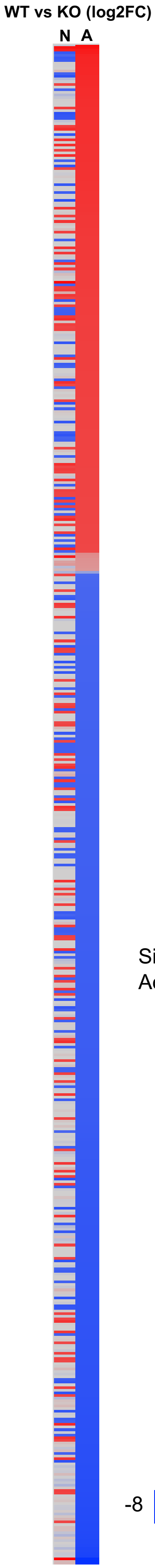


B.

| Ingenuity Canonical Pathways | -log(p-value) | Molecules |
|---|---|---|
| Systemic Lupus Erythematosus Signaling | 5.56 | CD247,CD2BP2,CD3G,CD86,HLA-A,HNRNPC,LAT,LSM14A,NFATC2,PRPF38B,PRPF40A,PRPF4B,PTPN6,PTPRC,SART1 |
| Calcium-induced T Lymphocyte Apoptosis | 4.17 | CD247,CD3G,HDAC1,HLA-A,ITPR2,NFATC2,PRKD3 |
| CD28 Signaling in T Helper Cells | 4.01 | CD247,CD3G,CD86,HLA-A,ITPR2,LAT,NFATC2,PTPN6,PTPRC |
| DNA Methylation and Transcriptional Repression Signaling | 3.74 | ARID4B,CHD4,HDAC1,MECP2,SUDS3 |
| Cdc42 Signaling | 3.58 | ARHGEF6,CD247,CD3G,CFL1,CLIP1,EXOC1,GSK3B,HLA-A,LLGL1,MYL12A |
| Nur77 Signaling in T Lymphocytes | 3.55 | CD247,CD3G,CD86,HDAC1,HLA-A,MAP3K3 |
| iCOS-iCOSL Signaling in T Helper Cells | 3.5 | CD247,CD3G,CD86,HLA-A,ITPR2,LAT,NFATC2,PTPRC |
| CTLA4 Signaling in Cytotoxic T Lymphocytes | 3.36 | CD247,CD3G,CD86,HLA-A,LAT,PPP2R5E,PTPN6 |
| Phospholipase C Signaling | 3.19 | ARHGEF18,ARHGEF6,CD247,CD3G,HDAC1,ITPR2,LAT,MYL12A,NFATC2,PEBP1,PRKD3,RPS6KA3 |
| Role of BRCA1 in DNA Damage Response | 2.84 | ARID1A,MSH6,PBRM1,RBL2,SMARCA4,SMARCC2 |
| Hereditary Breast Cancer Signaling | 2.84 | ARID1A,HDAC1,MSH6,PBRM1,POLR2A,SMARCA4,SMARCC2,XPC |
| B Cell Development | 2.66 | CD86,HLA-A,PTPRC,SPN |
| ATM Signaling | 2.42 | BRAT1,CBX3,PPP2R5E,TP53BP1,TRIM28,USP7 |
| Growth Hormone Signaling | 2.33 | PRKD3,PTPN6,RPS6KA1,RPS6KA3,SRF |
| Sumoylation Pathway | 2.29 | DAXX,HDAC1,KDM1A,PML,SENP7,SP1 |
| NER Pathway | 2.29 | ERCC4,HMGN1,POLR2A,TOP2B,USP7,XPC |
| T Cell Receptor Signaling | 2.25 | CD247,CD3G,LAT,NFATC2,PAG1,PTPRC |
| PD-1, PD-L1 cancer immunotherapy pathway | 2.23 | CBLB,CD247,GSK3B,HLA-A,LAT,PDCD4 |
| Role of NFAT in Regulation of the Immune Response | 2.16 | CD247,CD3G,CD86,GSK3B,HLA-A,ITPR2,LAT,NFATC2 |
| B Cell Receptor Signaling | 2.1 | APBB1IP,CFL1,GSK3B,MAP3K3,NFATC2,PAG1,PTPN6,PTPRC |
| Superpathway of Cholesterol Biosynthesis | 2.01 | ACAA2,DHCR24,LBR |
| PKCθ Signaling in T Lymphocytes | 2 | CD247,CD3G,CD86,HLA-A,LAT,MAP3K3,NFATC2 |
| 3-phosphoinositide Degradation | 2 | MTMR1,MTMR2,PPM1H,PPP1R8,PPP2R5E,PTPN6,PTPRC |

C.

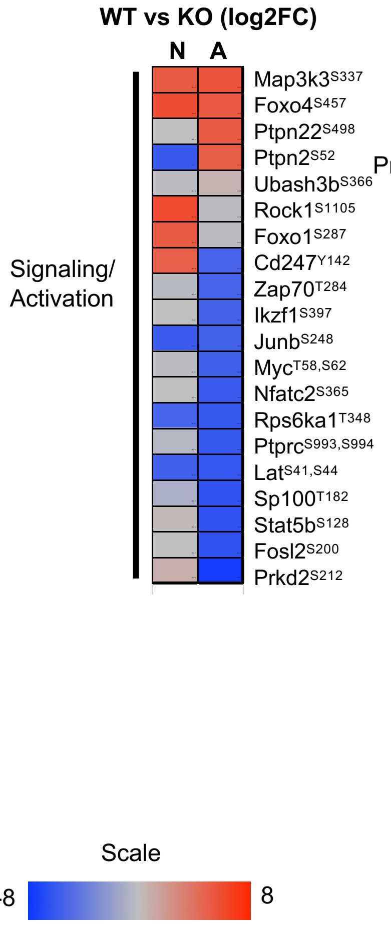


D.

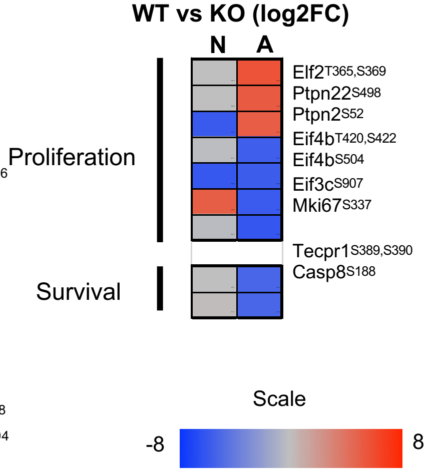


E.

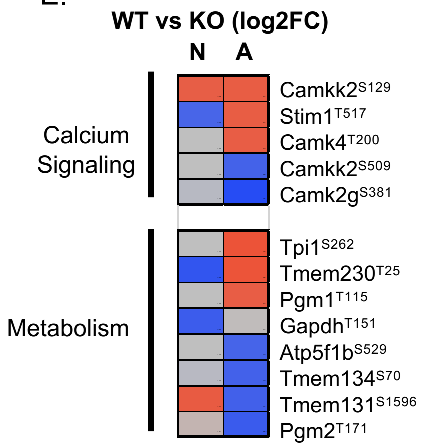


F.

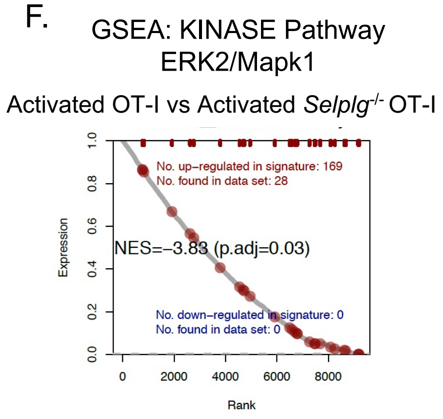

**Extended Data Figure 4.**

A.

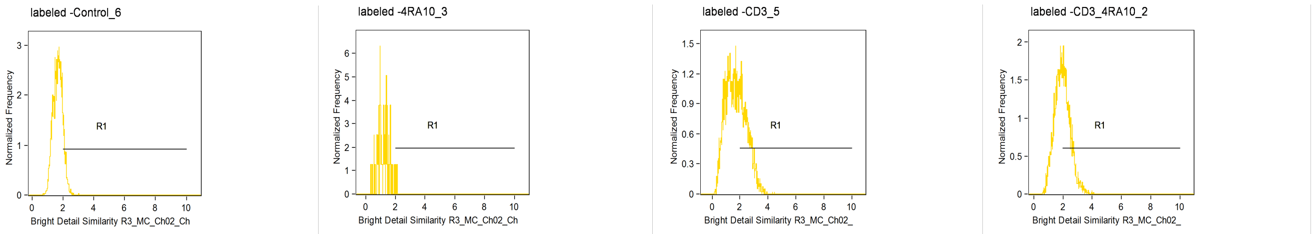


Bright Detail Similarity R3_MC_Ch02_Ch

| Population | Count | %Gated | Median | MAD |
|---|---|---|---|---|
| labeled & single cells & c... | 4452 | 100 | 1.667 | 0.3217 |
| R1 & labeled & single cell... | 571 | 12.8 | 2.118 | 0.1068 |

Bright Detail Similarity R3_MC_Ch02_Ch

| Population | Count | %Gated | Median | MAD |
|---|---|---|---|---|
| labeled & single cells & c... | 79 | 100 | 1.206 | 0.382 |
| R1 & labeled & single cell... | 2 | 2.53 | 2.083 | 0.04987 |

Bright Detail Similarity R3_MC_Ch02_

| Population | Count | %Gated | Median | MAD |
|---|---|---|---|---|
| labeled & single cells & c... | 3922 | 100 | 1.677 | 0.784 |
| R1 & labeled & single cell... | 1362 | 34.7 | 2.443 | 0.3985 |

Bright Detail Similarity R3_MC_Ch02_

| Population | Count | %Gated | Median | MAD |
|---|---|---|---|---|
| labeled & single cells & c... | 4457 | 100 | 1.94 | 0.5081 |
| R1 & labeled & single cell... | 2004 | 45 | 2.336 | 0.3016 |

B.

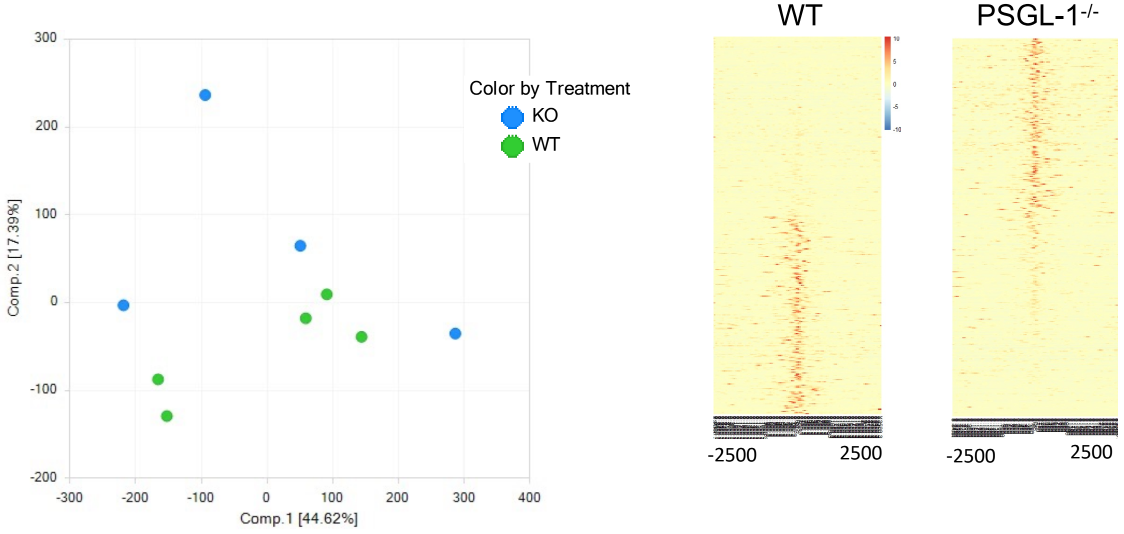


C.

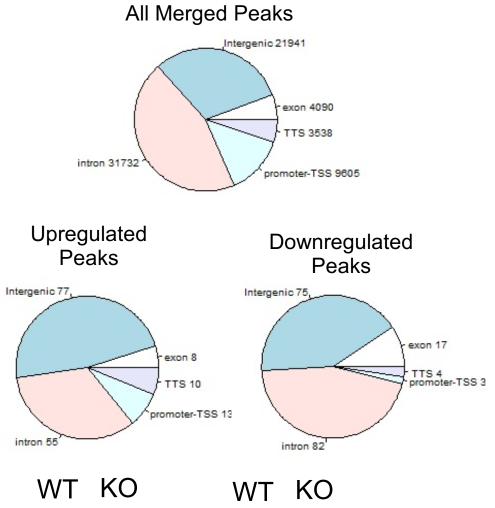


D.

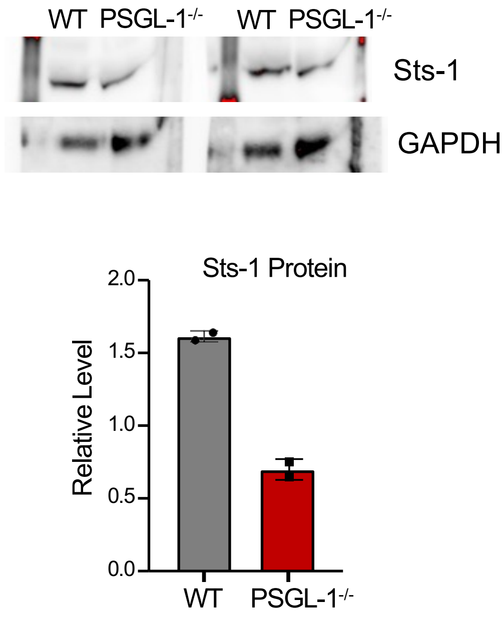

#### Extended Data Figure 5.

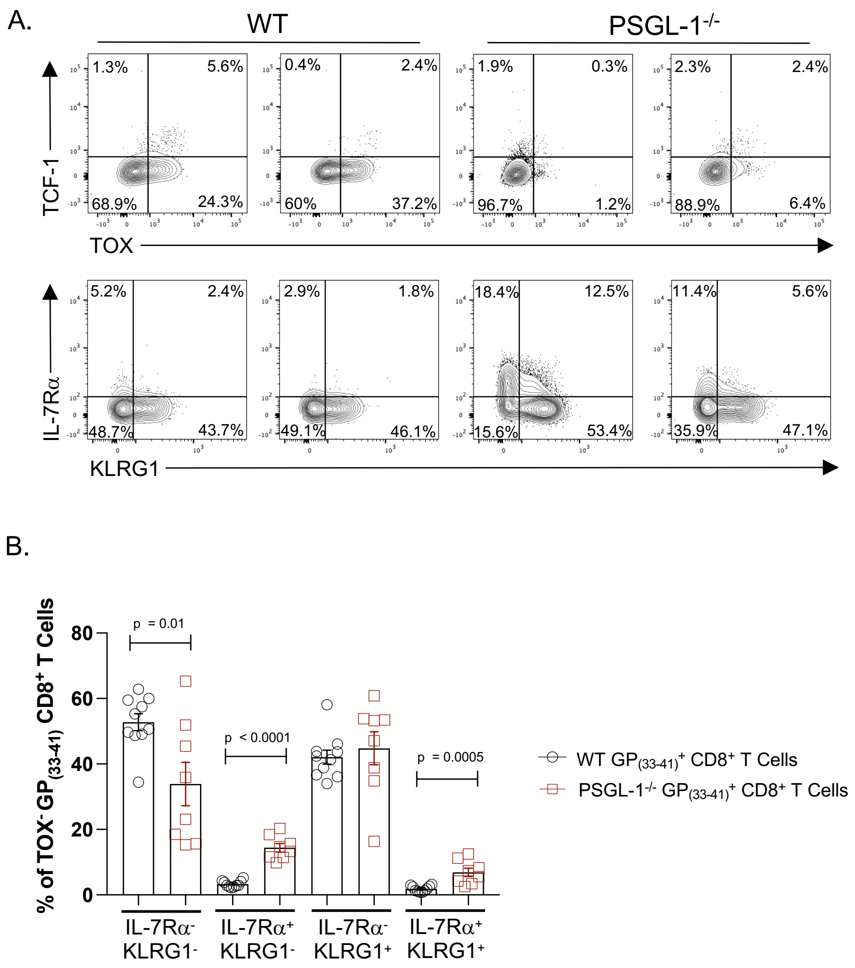

**Extended Data Figure 6.**

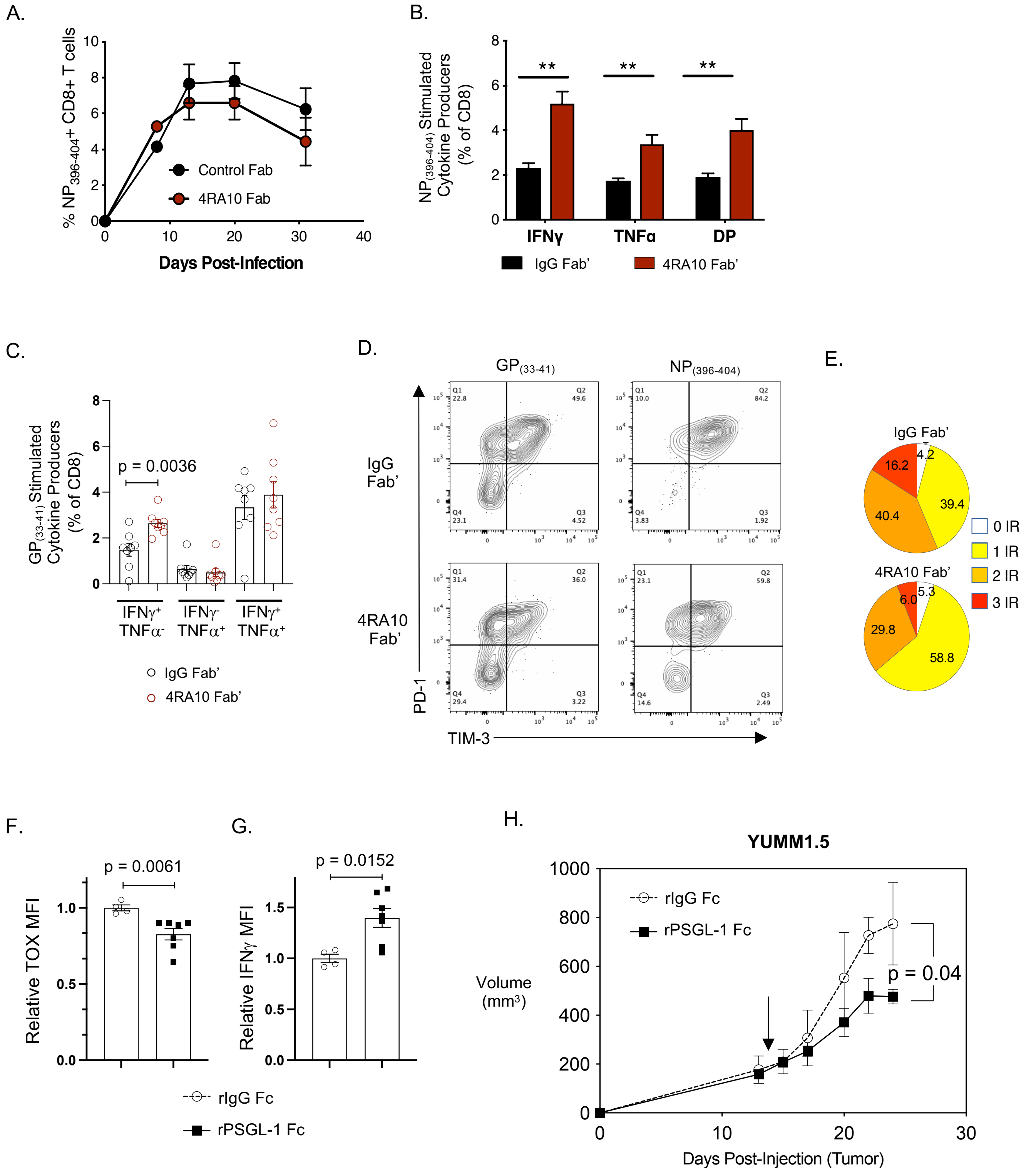

### Extended Data Table 1

| VENDOR | CAT# | ANTIBODY | CLONE | MIX |
|---|---|---|---|---|
| BD Biosciences | 563550 | BUV395 Mouse Anti-Human CD4 | SK3 | Surface Staining Flow Cytometry |
| BD Biosciences | 563786 | BUV395 Rat Anti-Mouse CD8a | 53-6.7 | Surface Staining Flow Cytometry |
| BD Biosciences | 612942 | BUV496 Mouse Anti-Human CD8 | RPA-T8 | Surface Staining Flow Cytometry |
| BD Biosciences | 612846 | BUV737 Mouse Anti-Human CD45RA | HI100 | Surface Staining Flow Cytometry |
| BD Biosciences | 612811 | BUV737 Mouse Anti-Mouse CD45.1 | A20 | Surface Staining Flow Cytometry |
| BD Biosciences | 741727 | BUV737 Rat Anti-Mouse CD11a | M17/4 | Surface Staining Flow Cytometry |
| BD Biosciences | 741820 | BUV737 Rat Anti-Mouse CD223 | C9B7W | Surface Staining Flow Cytometry |
| BD Biosciences | 740015 | BV421 Mouse Anti-Mouse CD244.2 | 2B4 | Surface Staining Flow Cytometry |
| BD Biosciences | 744284 | BV421 Rat Anti-Mouse CD244.1 | C9.1 | Surface Staining Flow Cytometry |
| BD Biosciences | 746479 | BV480 Mouse Anti-Human CD44 | L178 | Surface Staining Flow Cytometry |
| BD Biosciences | 745250 | BV605 Mouse Anti-Mouse Ly-108 | 1363 | Surface Staining Flow Cytometry |
| BD Biosciences | 751861 | R718 Rat Anti-Human CCR7 (CD197) | 3D12 | Surface Staining Flow Cytometry |
| BD Biosciences | 566939 | R718 Rat Anti-Mouse CD4 | RM4-5 | Surface Staining Flow Cytometry |
| BioLegend | 331010 | Alexa Fluor® 647 anti-human CD101 (BB27) | BB27 | Surface Staining Flow Cytometry |
| BioLegend | 304250 | APC/Fire™ 750 anti-human CD45RO | UCHL1 | Surface Staining Flow Cytometry |
| BioLegend | 102738 | APC/Fire™ 750 anti-mouse CD38 | 90 | Surface Staining Flow Cytometry |
| BioLegend | 427403 | Apotracker™ Green | N/A | Surface Staining Flow Cytometry |
| BioLegend | 303526 | Brilliant Violet 421™ anti-human CD38 | HIT2 | Surface Staining Flow Cytometry |
| BioLegend | 302632 | Brilliant Violet 605™ anti-human CD25 | BC96 | Surface Staining Flow Cytometry |
| BioLegend | 202537 | Brilliant Violet 605™ anti-rat CD90/mouse CD90.1 (Thy-1.1) | OX-7 | Surface Staining Flow Cytometry |
| BioLegend | 345024 | Brilliant Violet 711™ anti-human CD366 (Tim-3) | F38-2E2 | Surface Staining Flow Cytometry |
| BioLegend | 119727 | Brilliant Violet 711™ anti-mouse CD366 (Tim-3) | RMT3-23 | Surface Staining Flow Cytometry |
| BioLegend | 369322 | Brilliant Violet 785™ anti-human CD223 (LAG-3) | 11C3C65 | Surface Staining Flow Cytometry |
| BioLegend | 135225 | Brilliant Violet 785™ anti-mouse CD279 (PD-1) | 29F.1A12 | Surface Staining Flow Cytometry |
| BioLegend | 110706 | FITC anti-mouse CD45.1 | A20 | Surface Staining Flow Cytometry |
| BioLegend | 103606 | FITC anti-mouse CD49d | R1-2 | Surface Staining Flow Cytometry |
| BioLegend | 422302 | Human TruStain FcX™ | N/A | Surface Staining Flow Cytometry |
| BioLegend | 328208 | PE anti-human CD39 | A1 | Surface Staining Flow Cytometry |
| BioLegend | 143004 | PE anti-mouse CD160 | 7H1 | Surface Staining Flow Cytometry |
| BioLegend | 341212 | PE/Cyanine7 anti-human CD160 | BY55 | Surface Staining Flow Cytometry |
| BioLegend | 367434 | PE/Dazzle™ 594 anti-human CD279 (PD-1) | NAT105 | Surface Staining Flow Cytometry |
| BioLegend | 104448 | PE/Dazzle™ 594 anti-mouse CD62L | MEL-14 | Surface Staining Flow Cytometry |
| BioLegend | 140329 | PE/Dazzle™ 594 anti-mouse CD90.2 (Thy-1.2) | 53-2.1 | Surface Staining Flow Cytometry |
| BioLegend | 310926 | PerCP/Cyanine5.5 anti-human CD69 | FN50 | Surface Staining Flow Cytometry |
| BioLegend | 121626 | PerCP/Cyanine5.5 anti-mouse CD107a (LAMP-1) | 1D4B | Surface Staining Flow Cytometry |
| BioLegend | 125212 | PerCP/Cyanine5.5 anti-mouse CD223 (LAG-3) | C9B7W | Surface Staining Flow Cytometry |
| BioLegend | 101302 | Purified anti-mouse CD16/32 | 93 | Surface Staining Flow Cytometry |
| eBioscience/ThermoFisher | 25-1011-82 | CD101 PECy7 | Moushi101 | Surface Staining Flow Cytometry |
| NIH Tetramer Core | N/A | D(b) GP(33-41) APC or BV421 Tetramer | N/A | Surface Staining Flow Cytometry |
| NIH Tetramer Core | N/A | D(b) NP(396-401) BV421 Tetramer | N/A | Surface Staining Flow Cytometry |
| BD Biosciences | 557817 | Alexa Fluor 647 Mouse Anti-ZAP70 (PY319)/Syk (PY352) | 17A/P-ZAP70 | Intracellular Flow Cytometry |
| BD Biosciences | 564217 | PE Mouse Anti-TCF-7/TCF-1 | S33-966 | Intracellular Flow Cytometry |
| BD Biosciences | 564217 | PE Mouse Anti-TCF-7/TCF-1 | S33-966 | Intracellular Flow Cytometry |
| BioLegend | 502912 | APC anti-human TNF-α | MAb11 | Intracellular Flow Cytometry |
| BioLegend | 505810 | APC anti-mouse IFN-γ | XMG1.2 | Intracellular Flow Cytometry |
| BioLegend | 154304 | APC anti-mouse Perforin | S16009A | Intracellular Flow Cytometry |
| BioLegend | 502532 | Brilliant Violet 421™ anti-human IFN-γ | 4S.B3 | Intracellular Flow Cytometry |
| BioLegend | 396414 | Brilliant Violet 421™ anti-human/mouse Granzyme B | QA18A28 | Intracellular Flow Cytometry |
| BioLegend | 644816 | Brilliant Violet 421™ anti-T-bet Antibody | 4B10 | Intracellular Flow Cytometry |
| BioLegend | 500307 | PE anti-human IL-2 | MQ1-17H12 | Intracellular Flow Cytometry |
| BioLegend | 503808 | PE anti-mouse IL-2 | JES6-5H3 | Intracellular Flow Cytometry |
| BioLegend | 506324 | PE/Cyanine7 anti-mouse TNF-α | MP6-XT22 | Intracellular Flow Cytometry |
| eBioscience/ThermoFisher | 25-4875-82 | EOMES PECy7 | Dan11mag | Intracellular Flow Cytometry |
| eBioscience/ThermoFisher | 46-4877-42 | EOMES PerCP-eFluor710 | WD1928 | Intracellular Flow Cytometry |
| eBioscience/ThermoFisher | 46-5698-82 | Ki67 PerCP-eFluor710 | SolA15 | Intracellular Flow Cytometry |
| eBioscience/ThermoFisher | 48-5825-82 | T-bet eFluor 450 | eBio4B10 | Intracellular Flow Cytometry |
| eBioscience/ThermoFisher | 50-6502-82 | TOX eFluor 660 | TXRX10 | Intracellular Flow Cytometry |

**Extended Information Titles and Legends**

**Extended Data Figure 1. PSGL-1 restrains TCR stimulation strength and TCR signaling in T cells.** (A) Representative flow cytometry histograms of the data presented in Figure 1A. (B) Polyclonal C57BL/6 naïve $CD4^+$ and $CD8^+$ T cells isolated from C57BL/6 or PSGL-$1^{-/-}$ mice were stimulated with plate-bound anti-CD3ε antibody at the indicated concentration for 3 days prior to supernatant collection. Luminex cytokine assays were performed to quantify IL-2, IFNγ, and TNFα production. Data are representative of experimental duplicate wells from one experiment. (C) Western blot detection of phosphorylated and total levels of Zap70, Erk1/2, AKT and GAPHD in OT-I WT and PSGL-$1^{-/-}$ $CD8^+$ T cells stimulated for the indicated time with Streptavidin-conjugated anti-CD3ε antibody and crosslinking by anti-Strepavidin. Data are representative of three independent experiments.

**Extended Data Figure 2. PSGL-1 deficiency limits the development of T cell exhaustion in response to the lower affinity peptide SIIQFEKL.** Dot plots of the frequency of (A) IFNγ and TNFα double-producing-, (C) IFNγ-producing, and (E) TNFα-producing $CD44^+$ OT-I $CD8^+$ T cells cultured under $iT_{EFF}$ or $iT_{EX}$ conditions with either SIINFEKL or SIIQFEKL peptide for 5 days prior to 5 hour restimulation with SIINFEKL peptide. Lines indicate the matched WT OT-I and PSGL-$1^{-/-}$ OT-I $CD8^+$ T cells from the same experiment. Dot plots of the PSGL-$1^{-/-}$ OT-I to WT OT-I (KO/WT) ratio of (B) IFNγ and TNFα double-producing-, (D) IFNγ-producing, and (F) TNFα-producing $CD44^+$ OT-I $CD8^+$ T cells.

**Extended Data Figure 3. Phospho-proteomics analysis reveals increased expression of molecules associated with T cell activation in PSGL-1 deficient $CD8^+$ T cells.** Naïve WT OT-I and PSGL-$1^{-/-}$ OT-I $CD8^+$ T cells were incubated with immobilized isotype control IgG (naïve) or anti-CD3 antibodies for 15 minutes prior to collection and processing for proteomics and phosphoproteomics analysis. (A) Heatmap of 576 phosphorylated proteins differentially expressed ($\geq$ 2 log2FC, FDR $<$ 0.01) between OT-I and PSGL-$1^{-/-}$ OT-I $CD8^+$ T cells after 15 minutes of activation (A, right) and the coordinate expression in non-activated, naïve cells (N, left). Data are displayed as expression in WT OT-I (WT) vs PSGL-$1^{-/-}$ OT-I (KO) $CD8^+$ T cells. (B) List of the 22 canoncial pathways identified by IPA analysis with expression patterns potentially associated with activated WT OT-I vs PSGL-$1^{-/-}$ OT-I $CD8^+$ T cells. Heatmaps of selected genes and their expression in WT OT-I vs PSGL-$1^{-/-}$ OT-I $CD8^+$ T cells at the naïve (N) state and after 15 minutes of activation in association with (C) T cell signaling/activation, (D) proliferation and survival, and (C) calcium signaling and metabolism. (F) GSEA kinase pathway analysis of ERK2/Mapk1 in activated WT OT-I and PSGL-$1^{-/-}$ OT-I $CD8^+$ T cells.

**Extended Figure 4. *Ubash3b*/Sts-1 is downregulated in the absence of PSGL-1, promoting greater TCR signaling in $CD8^+$ T cells.** (A) Left: PCA plot of individual WT (green) and KO (blue) ATAC-seq libraries. Right: ATAC-seq profile of reproducible chromatin accessibility in WT and KO libraries. (B) Pie charts showing the breakdown by type in all, upregulated, and downregulated peaks. (C) Western blot of Sts-1 and GAPDH protein expression in WT OT-I and PSGL-$1^{-/-}$ OT-I $CD8^+$ T cells activated for 2 days with immobilized anti-CD3 and anti-CD28 (each 5 μg/mL). (D) Representative histograms from the Amnis ImageStream showing how the % positive similarity score was determined.

**Extended Data Figure 5. Increased frequency of IL-7Rα- and KLRG-1-expressing cells in $TOX^-$ PSGL-$1^{-/-}$ $CD8^+$ T cells during early LCMV Cl13 infection.** C57BL/6 WT or PSGL-$1^{-/-}$ mice were infected with LCMV Cl13 virus. Virus-specific T cell responses were assessed using $GP_{(33\text{-}41)}$ and $NP_{(396\text{-}404)}$ MHC-I tetramers in combination with flow cytometry. (A) Top: representative flow cytometry plots showing TCF-1 and TOX expression in $GP_{(33\text{-}41)}$-specific $CD8^+$ T cells from the blood of WT or PSGL-$1^{-/-}$ mice on day 9 post-infection. Bottom: representative flow cytometry plots showing IL-7Rα and KLRG1 expression in $TOX^-$ $GP_{(33\text{-}41)}$-specific $CD8^+$ T cells from the blood of WT or PSGL-$1^{-/-}$ mice on day 9 post-infection. (B) Dot plot/bar graphs of the frequency of IL-7Rα and KLRG1 expression in $GP_{(33\text{-}41)}^+$ $CD8^+$ T cells in the blood of WT or PSGL-$1^{-/-}$ mice on day 9 post-infection. Data are normally distributed (Shapiro-Wilk); unpaired t test was used for statistical analysis. Data is representative of one of two independent experiments. Each dot represents an individual mouse.

**Extended Data Figure 6. Fab-mediated inhibition of PSGL-1 promotes decreased T cell exhaustion and functional T cell responses to LCMV Cl13.** C57BL/6 mice were infected with LCMV Cl13 and treated I.P. with 100 μg of Fab' control rat IgG (control Fab) or Fab' anti-PSGL-1 (4RA10 Fab) in the AM and PM beginning on day 4 after LCMV Cl13 infection and subsequently on days 6, 8, 10, and 12 post-infection. Virus-specific T cells in the blood were assessed by flow cytometry on days 8, 13, 20, and 31 post-infection. (A) Kinetics of the frequency of $NP_{(396\text{-}404)}$-specific $CD8^+$ T cells in the blood of LCMV Cl13 infected C57BL/6 mice treated with either control Fab' or 4RA10 Fab'. (B) Bar graph of cytokine producing $CD8^+$ T cells on day 8 post-infection control Fab'- or 4RA10 Fab'-treated mice following restimulation with $NP_{(396\text{-}404)}$ peptide for 5 hours in the presence of brefeldin A. (C) Dot plot/Bar graph of cytokine producing $CD8^+$ T cells on day 8 post-infection control Fab'- or 4RA10 Fab'-treated mice following restimulation with $GP_{(33\text{-}41)}$ peptide for 5 hours in the presence of brefeldin A. (D) Representative FACS plots of PD-1 and TIM-3 expression in $GP_{(33\text{-}41)}$ (left) and $NP_{(396\text{-}401)}$ (right) virus-specific $CD8^+$ T cells in mice treated with control Fab' (top) or 4RA10 Fab' (bottom) at day 8 post-infection with LCMV Cl13. (E) Pie graphs of combinatorial expression of PD-1, LAG3, and TIM-3 on $NP_{(396\text{-}404)}$-specific $CD8^+$ T cells on day 31 post-infection in control Fab'- or 4RA10 Fab'-treated mice. Data assessed via Boolean gating. (F) Relative TOX expression (MFI) in $TOX^+$ adoptively transferred WT P14 ($CD45.1^+$) $CD8^+$ T cells in C57BL/6 ($CD45.2^+$) mice infected with LCMV Cl13 and treated with control Fc (rIgG Fc) or recombinant PSGL-1 (rPSGL-1 Fc) protein on days 0, 3, and 6 post-infection. (G) Relative IFNγ expression (MFI) in $IFN\gamma^+$ adoptively transferred WT P14 ($CD45.1^+$) $CD8^+$ T cells as in F, following overnight restimulation with $GP_{(33\text{-}41)}$ peptide in the presence of Brefeldin A. (H) Average tumor growth (volume) in YUMM1.5 tumor-bearing mice treated with either control Fc or rPSGL-1 Fc protein with treatment initiated on day 14 post-injection of YUMM1.5 tumor cells.
